# SilkRoute: A Descriptor-Driven Framework for Reproducible Multi-Source Biomolecular Data Acquisition

**DOI:** 10.64898/2026.08.11.744100

**Authors:** Diego Fernández, Julián García-Vinuesa, Diego Alvarez-Saravia, Michelle Soto-García, José L. Medina-Franco, Julieta Sepulveda-Yanez, Xavier Cadet, Frederic Cadet, Mehdi D. Davari, Roberto Uribe-Paredes, Fabio Herrera-Rocha, David Medina-Ortiz

## Abstract

**Background:** Biomolecular dataset construction often requires coordinated retrieval from heterogeneous repositories, identifier mapping, cross-reference enrichment, source-specific parsing, and provenance recording. These operations are frequently implemented through project-specific scripts, making acquisition procedures difficult to inspect, reproduce, or adapt across studies. We present SilkRoute, an open-source Python framework that formalizes biomolecular data acquisition as descriptor-defined, source-aware, and provenance-tracked workflows, providing a reproducible foundation for multi-source biomolecular dataset construction.

**Results:** SilkRoute uses machine-readable YAML descriptors to specify dataset intent, biomolecular modality, workflow mode, query logic, enrichment resources, execution parameters, and export settings. These descriptors drive a common execution model that coordinates primary retrieval and downstream enrichment while preserving source-specific outputs, interaction evidence when available, the original workflow configuration, metadata, and run summaries. We evaluated this model through three representative acquisition scenarios spanning proteins, compounds, and molecular interactions. In the protein-centered workflow, SilkRoute retrieved 2,444 reviewed antimicrobial protein records from UniProt and generated complementary outputs from AlphaFold DB, InterPro, Pathway Commons, and the Protein Data Bank. In the compound-centered workflow, a ChEMBL IC_50_ query produced 1,445,939 activity records organized into query-defined potency ranges. In the interaction-centered workflow, 2,253 UniProt protein records were expanded with 902,713 BioGRID interaction records and 5,702 STRING interaction-partner records. Across these scenarios, the framework successfully applied the same descriptor-defined acquisition model to distinct biomolecular entity types, retrieval strategies, enrichment paths, and output structures.

**Conclusions:** SilkRoute extends beyond sequence retrieval by providing a reusable acquisition layer for constructing multi-source biomolecular datasets. By separating primary retrieval from enrichment and preserving source-aware outputs together with workflow descriptors and execution metadata, the framework makes acquisition procedures easier to inspect, reproduce, archive, and adapt. SilkRoute does not replace biological curation, label validation, deduplication, partitioning, or benchmarking, but provides structured and traceable acquisition packages that support these downstream processes.

## 1 Background

Computational biology increasingly relies on datasets that combine molecular entities, annotations, measurements, and contextual metadata distributed across multiple biological repositories [1]. This dependence is particularly evident in biomolecular machine-learning applications, including protein function prediction [2], protein engineering [3], therapeutic molecule discovery (e.g., peptides) [4, 5], molecular interaction modeling [6], and bioactivity prediction [7]. Such applications commonly require the joint use of protein sequences, structural models, experimental structures, functional domains, pathway annotations, interaction evidence, biochemical properties, compound records, and bioactivity measurements [8].

For studies built from distributed biomolecular evidence, dataset construction is not a secondary technical step [9]. The acquisition process determines which biological entities are represented, which annotations are associated with them, which repositories are treated as authoritative, and which assumptions are propagated into downstream analyses [10]. Consequently, the quality and traceability of acquisition workflows directly influence the reliability of computational studies, together with model design, statistical evaluation, and biological interpretation [11].

Despite the maturity of major biological databases, the information required to construct biomolecular datasets remains distributed across resources with different scopes, identifiers, metadata, query systems, access policies, update cycles, and data formats [12, 13, 9]. Protein-centered workflows may combine sequence data from UniProt [14]; structural information from the Protein Data Bank [15] or AlphaFold DB [16]; functional classifications from Gene Ontology [17]; pathway knowledge from Reactome [18] or Rhea [19]; and interaction evidence from STRING [20] and BioGRID [21]. Multi-omics studies introduce further resources, including ChEMBL [22], PubChem [23], ChEBI [24], BRENDA [25], SABIO-RK [26], and PRIDE [27], which organize chemical entities, bioactivities, enzyme properties, kinetic measurements, and proteomics evidence.

The challenge therefore arises not from the absence of curated data sources, but from the need to coordinate them within a single acquisition process. Constructing a reusable biomolecular dataset commonly requires coordinated queries, identifier mapping, cross-reference resolution, source-specific parsing, field selection, metadata integration, and export decisions [9]. The central difficulty is the reconstruction of the computational path that connects a biological question to a structured dataset. In practice, this path is frequently encoded in project-specific scripts, notebooks, command histories, or manually documented procedures that are difficult to inspect, rerun, compare, or adapt across studies.

This limitation becomes particularly evident when acquisition is treated as an informal preprocessing step. A dataset may be described through its final table, while the procedures used to retrieve and enrich the data remain only partially documented [28]. Such opacity limits reproducibility and reuse. Changes in database structure and policies can alter retrieved content even when the same biological query is repeated [29, 30]. Without explicit acquisition protocols and provenance records, it becomes difficult to determine the source of differences between dataset results. These concerns are consistent with broader challenges in computational reproducibility, where executable workflows, provenance capture, and transparent data-processing histories are essential for making analyses auditable and reusable [31, 32, 11, 33].

Existing software ecosystems provide important components for biological data access. Biopython offers broad programmatic utilities for sequence analysis, file parsing, and interaction with biological resources [34], while BioServices provides unified Python access to multiple biological web services and databases [35]. Repository-specific APIs additionally enable direct access to resources such as UniProt, PDB, ChEMBL, PubChem, Reactome, KEGG, STRING, and BioGRID, whereas identifier-mapping resources support cross-database navigation and annotation retrieval [13].

Complementing these interfaces, workflow systems such as Galaxy, CWL-based platforms, and Nextflow provide reproducible execution, portability, dependency management, and provenance for computational analyses [36, 37, 38]. Data-recipe approaches likewise support the preservation and reuse of data-processing procedures [39]. Although these platforms effectively coordinate computational workflows, the acquisition stage remains largely repository specific. Users must therefore define or integrate data sources, query logic, and output conventions for each biomolecular dataset.

Consequently, biomolecular data acquisition is often implemented through custom scripts or workflow definitions that may differ substantially across projects. While these approaches are flexible, they are typically weakly standardized, difficult to maintain, and rarely capture acquisition intent, repository relationships, execution context and provenance in a unified reusable form. Existing tools therefore remain foundational for biological data retrieval but generally do not offer a domain-specific solution for biomolecular data acquisition workflows at the dataset level.

A domain-specific acquisition layer can bridge this gap by connecting biological repositories with downstream analysis frameworks [40]. Rather than replacing established repositories, database clients, and curation standards, such a layer formalizes how biological questions are translated into executable acquisition protocols, how primary records are retrieved and enriched through external identifiers, how heterogeneous outputs are organized, and how provenance is recorded [31]. Importantly, this layer remains distinct from downstream processes including biological curation, label validation, leakage control, dataset partitioning, representation learning, and model benchmarking [41, 42]. Its purpose is to produce structured, traceable and reproducible datasets that can subsequently be inspected, archived, curated, transformed, and reused in bioinformatics and machine-learning workflows [43].

Here, we present SilkRoute v0.1.0, an open-source Python framework that formalizes biomolecular data acquisition as a reproducible and traceable foundation for dataset construction from heterogeneous biological repositories. SilkRoute uses machine-readable YAML descriptors to capture dataset intent, biomolecular modality, workflow mode, query logic, enrichment configuration, execution parameters, export settings, and provenance requirements. It separates primary retrieval from downstream enrichment, allowing initial record sets to be extended with structural, functional, pathway, interaction, chemical, or biochemical information through source-specific interfaces and cross-references. The framework supports both command-line and programmatic execution, while an optional graphical descriptor builder simplifies the creation of executable YAML workflow configurations. These interfaces share a common acquisition model and execution components. By building on established repository APIs and Python bioinformatics libraries, SilkRoute provides a high-level abstraction for defining, executing, documenting, and reviewing the upstream acquisition processes involved in biomolecular dataset construction. We demonstrate its capabilities through representative protein-, compound-, and interaction-centered workflows, including the acquisition of reviewed antimicrobial protein, ChEMBL IC_50_ activity retrieval, and protein–protein interaction enrichment.

## 2 Implementation

### 2.1 Design principles and overall architecture

SilkRoute was developed as a modular Python framework for defining and executing biomolecular data acquisition workflows across heterogeneous biological repositories. Its implementation was guided by six complementary design principles intended to make acquisition procedures explicit, adaptable, and traceable while preserving the operational differences among external data sources.

A central design objective was to make acquisition procedures declarative. Dataset intent, biomolecular modality, workflow mode, query logic, enrichment configuration, execution parameters, and export settings are therefore specified before execution in machine-readable YAML descriptors. This separates workflow specification from the source code responsible for execution, allows acquisition protocols to be reviewed, versioned, reused, and archived independently of the implementation.

Separating workflow specification from execution naturally requires isolating repository-specific behavior from the general execution logic. Biological repositories differ in endpoint organization, accepted identifiers, query syntax, pagination mechanisms, authentication requirements, response structures, and field semantics. SilkRoute encapsulates these differences within dedicated source interfaces, allowing changes in an external service to be accommodated locally without requiring modifications throughout the workflow engine.

Once repository-specific communication has been isolated, the framework distinguishes between primary retrieval and enrichment. Primary retrieval defines the initial collection of records, whereas enrichment extends those records with structural, functional, pathway, interaction, chemical, or biochemical information obtained through external identifiers and source-specific interfaces. Treating these as distinct stages makes the origin and purpose of every retrieved component explicit.

Maintaining this separation also requires preserving the inherent heterogeneity of biological repositories. Protein records, interaction evidence, pathway payloads, structural annotations, and compound activity measurements differ substantially in structure and interpretation. Rather than forcing these heterogeneous data into a universal flattened schema, SilkRoute preserves them as source-aware outputs, retaining repository-specific semantics for downstream analysis.

Preserving source-specific information extends beyond the retrieved data to the execution process itself. Provenance information is therefore generated as an integral part of every workflow execution. In addition to the retrieved records, SilkRoute stores the workflow descriptor and records the normalized execution parameters, generated files, execution status, record counts, runtime information, and source-level metadata where available. This enables every generated dataset to be traced back to the acquisition protocol and execution configuration that produced it.

These principles also define the intended functional scope of the framework. SilkRoute focuses on upstream data acquisition, enrichment, structured export, and provenance tracking. Downstream tasks, including biological curation, final label validation, advanced deduplication, dataset partitioning, representation learning, and model benchmarking, intentionally remain outside its scope because they depend on the specific biological question and the intended analytical use of the acquired data.

The six principles directly informed the intended layered architecture of SilkRoute, which separates workflow specification, execution control, repository communication, response interpretation, enrichment, output generation, and provenance reporting (Table **1**, Section S1 of Supplementary Information). Rather than treating these responsibilities as a single execution pipeline, the framework assigns each to dedicated components that can evolve independently while preserving a consistent workflow execution model across heterogeneous repositories.

**Table 1:** Implementation layers, responsibilities, and technical challenges addressed in SilkRoute.

| Layer | Responsibility | Technical challenge addressed |
| --- | --- | --- |
| Descriptor handling | Loads and validates **workflow-v1** YAML descriptors and converts user-defined values into normalized workflow parameters. | Separates acquisition protocols from source-code modifications and detects invalid workflow structures before execution. |
| Workflow orchestration | Determines the execution path, initializes the required source interfaces, and coordinates primary retrieval and enrichment. | Provides a common control flow across biomolecular modalities and workflow organizations. |
| Source-specific interfaces | Encapsulate endpoint configuration, request construction, identifier handling, pagination, batching, authentication, retry behavior, caching when available, and source-level error handling. | Localizes repository-specific behavior and limits the impact of upstream API changes on the broader codebase. |
| Parsing and transformation | Convert source responses into internal Python objects, tabular records, or structured payloads. | Decouples response interpretation from network communication and output generation. |
| Enrichment modules | Use accessions, cross-references, gene symbols, organism identifiers, compound identifiers, target identifiers, or other source-specific values to retrieve complementary records. | Extends primary records without collapsing structurally and semantically different resources into a single schema. |
| Export layer | Writes primary records, enrichment tables, interaction evidence, and structured source payloads according to the workflow configuration. | Preserves heterogeneous, relational, or nested outputs that cannot be represented appropriately in a single flattened table. |
| Metadata and reporting | Records the original descriptor, normalized parameters, source configuration, execution status, generated files, record counts, runtime information, and available source-level metadata. | Links generated outputs to both the requested acquisition protocol and the observed execution context. |
| Access interfaces | Provide descriptor-based execution through the command-line interface, programmatic access through the Python API, and guided descriptor generation through the graphical builder. | Supports different interaction modes without duplicating the underlying acquisition logic. |

Descriptor handling provides the entry point by validating the workflow specification and converting userdefined values into a normalized internal representation. Repository-specific components perform primary retrieval and optional enrichment, while dedicated parsing and transformation routines convert source responses into internal records or structured payloads. Export and reporting components organize the resulting objects into source-aware files and generate the associated execution records.

These architectural layers are coordinated by the MainWorkflow component. Following descriptor validation and normalization, MainWorkflow receives the biomolecular modality, workflow mode, executable query, execution parameters, and enrichment configuration. It determines the appropriate execution path, initializes the required source-specific interfaces, and coordinates primary retrieval and enrichment according to the normalized workflow definition. The same orchestration core is used for both command-line and programmatic execution, ensuring that workflow resolution and repository communication remain consistent regardless of the selected access mode (Section S1 of Supplementary Information).

### 2.2 Descriptor model and workflow resolution

The descriptor model defines how an acquisition workflow is represented and interpreted by the descriptor-handling and orchestration layers summarized in Table **1**. SilkRoute uses versioned YAML workflow descriptors, and workflow-v1 denotes the first implemented version of this descriptor schema. This schema provides a compact representation of an acquisition protocol, complementing biological metadata standards and dataset documentation by recording how biomolecular records should be selected, expanded, exported, and documented. A workflow-v1 descriptor is organized into sections describing dataset-level information, query definition, execution parameters, optional harmonization fields, export behavior, and reporting requirements.

The structure and major components of the workflow-v1 descriptor are described in Section S3 of the Supplementary Information, while Section S5.1 provides a complete protein-centered workflow illustrating its practical use. In this example, the descriptor defines a query for reviewed antimicrobial proteins, specifies the protein modality and query_first workflow mode, selects the UniProt fields to be retrieved [44], configures cross-reference enrichment through AlphaFold DB [16], PDB [15], Pathway Commons [45], and InterPro [46], and records the corresponding execution and export parameters.

The descriptor separates the biological acquisition objective from the operational settings required to execute it. The dataset section records the dataset identity, biomolecular modality, and workflow mode. The query section contains the executable selection criteria, requested source fields, and enrichment resources. Runtime behavior is defined through the execution section, while the export section specifies the output directory, file format, and reporting files to be generated. The optional harmonization section records selected semantic column roles and reporting information without imposing a universal schema on source-specific records.

During descriptor handling, the YAML specification is checked for the required structure and converted into a normalized internal representation (Section S3 of Supplementary Information). This step resolves user-facing values, applies defaults where required, and verifies that the declared modality, workflow mode, query organization, and execution options can be interpreted by the framework. Normalization therefore separates the human-readable workflow specification from the internal parameters required by the orchestration and source-integration components.

The execution path is determined from the combination of biomolecular modality, workflow mode, and query definition rather than from a single source identifier. This design avoids relying on a descriptive source field as the sole routing mechanism. Instead, the complete workflow configuration determines which source-specific interfaces are initialized and how the acquisition task is organized. The normalized query and execution parameters are then passed to the corresponding interfaces for source-level request construction.

Two workflow modes are currently implemented. In query_first mode, a primary query defines the initial record set, which can subsequently be expanded through configured enrichment resources. In the representative protein-centered workflow, UniProt provides the primary protein records, while identifiers and cross-references associated with those records are used to retrieve complementary information from AlphaFold DB, PDB, Pathway Commons, and InterPro. Further details on the implemented query modes and executable query fields are provided in Section S3.1 of the Supplementary Information.

In query_composition mode, multiple query components define labeled acquisition subsets that are executed and combined within the same workflow. This mode can be used, for example, to retrieve ChEMBL activity records corresponding to different IC_50_ ranges. The resulting labels identify the query component through which each record was acquired and should therefore be interpreted as operational acquisition labels, not as biologically or pharmacologically validated classes; their interpretation and implementation in the ChEMBL IC50 query-composition workflow are further detailed in Sections S3.1 and S5.2.1–S5.2.2 of the Supplementary Information.

Source-specific interfaces translate normalized query definitions into the syntax and parameters required by the corresponding repository. Selected user-facing expressions can consequently be converted into explicit source-level constraints (section S3.1 of the Supplementary Information). For example, a compact UniProt sequence-length expression can be rewritten using the range syntax expected by the UniProt REST API, whereas a ChEMBL IC_50_ expression can be translated into constraints over standard_type, standard_value, and standard_units. These transformations are explicitly implemented for supported sources and query patterns. The interpretation layer does not infer unsupported biological concepts or function as a general ontology-reasoning system.

The original descriptor is preserved without modification in the generated output package. Normalized workflow values, resolved query parameters, source-level execution information, generated files, and execution outcomes are recorded separately by the metadata and reporting layer. The descriptor therefore functions both as an executable workflow specification and as a versionable record connecting the intended acquisition procedure with the resulting data package (section S4.2 of Supplementary Information).

### 2.3 Source integration and critical implementation issues

After workflow resolution, acquisition is delegated to source-specific interfaces that encapsulate the operational behavior of each external repository. These interfaces address differences in endpoints, accepted identifiers, query syntax, pagination, response formats, authentication requirements, rate limits, and field semantics. Each interface implements only the query builders, retrieval operations, parsers, and enrichment routines required for its supported workflow roles. Support for a repository therefore refers to explicitly implemented operations and does not imply exhaustive coverage of every endpoint or data type exposed by the upstream resource (section S2 of Supplementary Information).

Source-specific request construction separates repository syntax from the general workflow logic. The orchestration layer passes normalized acquisition parameters to the selected interface, which converts them into the request structure expected by the external service. This allows source-level query behavior to be modified without changing the descriptor model or the common orchestration components.

Large result sets and endpoint-specific pagination are also handled within the corresponding interfaces. Depending on the repository, requests can be divided into pages or batches, while concurrency and retry behavior can be controlled through execution parameters such as max_workers and total_retries. Individual interfaces may also use caching when supported. These mechanisms reduce unnecessary repeated requests and improve resilience to temporary network or service interruptions, although successful execution remains dependent on the availability and behavior of the external repositories.

Missing identifiers, incomplete cross-references, unavailable records, empty enrichment responses, and parsing failures are treated as acquisition outcomes rather than as evidence of biological absence. Interfaces handle these conditions according to the behavior of the corresponding service, and available status or failure information is propagated to the execution metadata and reporting outputs. SilkRoute does not infer missing biological annotations or convert unsuccessful retrievals into negative biological labels. The factors affecting enrichment coverage and the implications of incomplete source metadata for biological interpretation are further discussed in Sections S2 and S6.1 of the Supplementary Information.

Authentication is handled independently from the workflow descriptor. Public HTTP-based workflows can be executed without credentials, whereas selected services may require an API key, registered credentials, or a user email. These values are supplied through environment variables or local configuration files, including optional .env files, and are not intended to be stored in version-controlled workflow descriptors. This separation keeps acquisition protocols shareable while preventing private credentials from becoming part of the executable specification (section S2 of the Supplementary Information).

The modular interface design also limits the effects of upstream API evolution. Changes to endpoints, parameter names, response structures, or authentication requirements can be addressed within the affected source module without modifying the orchestration layer. This architecture does not remove the framework’s dependence on external services, but it localizes the maintenance required when those services change.

Cross-reference-based enrichment connects primary records with complementary resources. Depending on the workflow, protein accessions, database cross-references, gene symbols, organism identifiers, compound identifiers, or target identifiers can be passed to enrichment interfaces. Interaction-oriented sources are handled through dedicated routines because their outputs represent relationships between entities rather than additional attributes of individual primary records. See Sections S1 and S5.1.4 of the Supplementary Information for implementation details.

### 2.4 Output organization, provenance, and reproducibility

The export and reporting layers convert the records produced during acquisition into a source-aware output package. Primary records, enrichment results, interaction evidence, and structurally complex responses are written as separate outputs when their schema or biological interpretations differ. This avoids forcing protein records, structural annotations, pathway graphs, interaction edges, compound activities, and enzyme metadata into a single normalized table. Representative export structures and output schemas are described in Section S4.3 of the Supplementary Information.

When available, enrichment outputs retain contextual identifiers connecting the retrieved information with the corresponding primary records and external source. These identifiers are generated by the relevant source-specific interface and allow downstream users to determine how enrichment records relate to the initial acquisition set. SilkRoute does not automatically impose a universal cross-database schema or determine how heterogeneous outputs should be merged for a particular biological analysis.

The optional harmonization section supports a limited set of export and reporting operations. For example, id_column can be used to add deterministic row identifiers to exported tabular copies when the requested column is absent, while sequence_column can support reporting of unique non-null sequences. The label_column and metadata_fields entries are retained as descriptor information. These fields do not independently rename, merge, deduplicate, or biologically harmonize source-specific records. The behavior and current scope of these descriptor fields are detailed in Section S4.1 of the Supplementary Information.

Responses whose structure cannot be represented adequately in a table are exported as separate structured payloads. Graph-like or nested records can therefore be written as structured files, including compressed JSON files, while a tabular index or summary maintains the relationship between the originating query and the external payload. This strategy is used, for example, for Pathway Commons neighborhood responses, where raw graph content is retained independently from its tabular summary. Representative export schemas and the Pathway Commons graph-payload implementation are described in Sections S4.3 and S5.1.6 of the Supplementary Information.

The provenance model distinguishes the planned acquisition protocol from the observed execution. The YAML descriptor records the workflow requested by the user, including its query logic, modality, workflow mode, enrichment configuration, execution parameters, and export settings. Following execution, metadata.json stores a detailed machine-readable manifest containing the original descriptor, normalized workflow values, source-specific parameters, generated outputs, execution status, runtime information, and available source-level metadata. The run_summary.yml file provides a compact overview of the workflow configuration, execution status, generated files, and record counts.

This distinction is important because procedural reproducibility does not guarantee that an external database will return identical biological content at a later date. Database releases, annotations, identifiers, endpoints, access policies, and service availability can change between executions. SilkRoute records how an acquisition was performed and preserves the outputs obtained during the reported run. Exact reconstruction of historical biological content additionally depends on archived output packages, stable database releases, or source snapshots. Further details on the relationship between workflow descriptors, execution manifests, and reproducibility under evolving external data sources are provided in Sections S4.2 and S6.1 of the Supplementary Information.

### 2.5 Access modes, implementation environment, and deployment

SilkRoute provides three complementary access modes built on the same descriptor model and orchestration components. The command-line interface supports complete descriptor-defined execution, including descriptor loading, validation, acquisition, enrichment, output writing, and reporting. The Python API exposes the orchestration core for integration into scripts, notebooks, and larger computational workflows, returning retrieved records and execution metadata in memory. The optional graphical descriptor builder assists users in creating, loading, validating, and exporting workflow-v1 descriptors through form-based inputs. Further details on graphical descriptor generation and representative command-line and programmatic execution are provided in Sections S3.2 and S5.1.2–S5.1.3 of the Supplementary Information.

The graphical builder does not implement an independent acquisition engine. It uses the shared descriptor schema and validation rules to generate workflow files that can subsequently be executed through the command-line interface. Similarly, command-line acquisition is delegated to the same MainWorkflow orchestration core exposed through the Python API. This arrangement ensures consistent workflow interpretation across access modes while retaining responsibilities appropriate to each interface.

SilkRoute was developed and continuously tested under Linux and was additionally subjected to manual installation and workflow-execution tests under Windows. Cross-platform support required attention to command invocation, file-system paths, process execution, and platform-specific launch behavior. Platform-independent path and configuration handling were used wherever possible to preserve consistent workflow interpretation across the supported operating systems.

The same workflow-v1 descriptors were used during the manual Windows validation, including checks of package installation, command-line invocation, path handling, descriptor loading, primary retrieval, enrichment execution, and output generation.

Development and workflow testing were performed in isolated Python environments. The reference environment used Python 3.13.3, while the package metadata declares compatibility with Python >=3.11,<3.15. YAML descriptors are parsed using PyYAML [47], tabular processing and export are implemented using polars [48], and sequence-oriented operations can use Biopython when required by individual source interfaces [34]. Dependencies required by the optional NiceGUI descriptor builder are maintained separately from the core workflow-execution requirements.

Endpoint definitions, supported fields, and other source-level configurations required by standard workflows are distributed with the package and loaded during execution. Credential-dependent values remain external to the distributed workflow descriptors, as described above. Installation requirements, supported data services, and external service configuration are described in Section S2 of the Supplementary Information.

### 2.6 Functional validation strategy

SilkRoute was functionally evaluated through representative protein-, compound-, and interaction-centered workflows. These scenarios were selected to determine whether the same descriptor-defined architecture could support different biomolecular entity types, primary sources, query organizations, enrichment paths, and output structures. The evaluation focused on software behavior and acquisition traceability rather than on deriving new biological conclusions from the retrieved records. The workflow configurations and representative outputs used in this evaluation are described in Section S5 of the Supplementary Information.

In addition to the end-to-end workflows, SilkRoute was evaluated using an automated test suite covering descriptor validation, workflow routing, query interpretation, command-line behaviour, graphical descriptor handling, source-specific parsers, and core acquisition components. Continuous integration was used to execute the automated tests under Python 3.11, 3.12, 3.13, and 3.14 in the reference Linux environment.

Each representative workflow was assessed from descriptor loading to final output generation. The evaluation covered structural validation, query normalization, source-specific request construction, response parsing, primary retrieval, enrichment execution, output writing, metadata-manifest generation, and run-summary creation. Generated directories were inspected to determine whether the primary records, source-specific enrichment outputs, interaction files, structured payloads, metadata manifests, and execution summaries specified by the workflow were produced.

Descriptor-based reproducibility was assessed by executing each workflow directly from its YAML specification and preserving that specification together with the generated outputs. The normalized runtime values recorded in metadata.json were used to inspect how the workflow had been interpreted during execution.

Source integration was assessed using open and credential-dependent services where applicable, while missing identifiers, incomplete cross-references, empty responses, unavailable records, authentication requirements, and schema differences were considered potential execution conditions. Further details on the relationship between workflow descriptors, execution metadata, and the generated acquisition packages are provided in Sections S4.2 and S5 of the Supplementary Information.

Functional consistency across access modes was assessed using the command-line interface for complete descriptor execution, the Python API for in-memory acquisition, and the graphical builder for descriptor generation. This assessment examined whether the access modes relied on the same descriptor specification and execution components; it was not designed as a formal user study. Representative examples of descriptor generation, command-line execution, and programmatic execution are provided in Sections S3.2 and S5.1.2–S5.1.3 of the Supplementary Information. Workflow-specific outputs and quantitative findings are presented in the Results and Discussion section.

## 3 Results and Discussion

### 3.1 Functional coverage, supported resources, and user-facing access

SilkRoute v0.1.0 was functionally evaluated across protein-, compound-, and interaction-centered acquisition scenarios. In each case, the workflow was initiated from a machine-readable specification and completed through primary retrieval, optional source-specific expansion, output generation, and execution reporting. The resulting runs produced primary records, enrichment or interaction outputs when configured, and provenance files connecting the generated artifacts with their acquisition context (Figure **1**).

**Figure 1:**
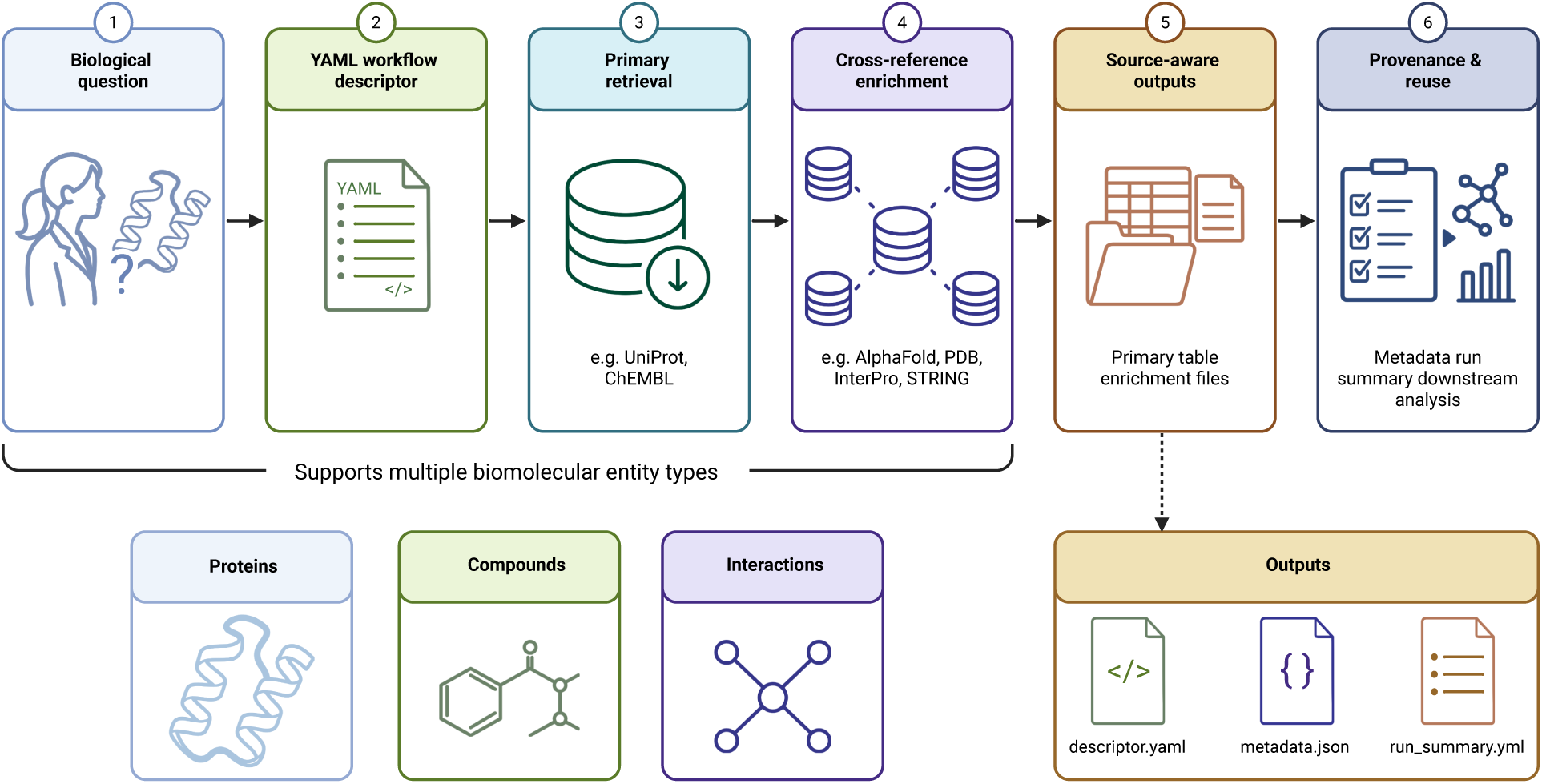
Descriptor-driven biomolecular data acquisition with SilkRoute. A biological acquisition objective is represented as a machine-readable YAML workflow descriptor that specifies the biomolecular modality, query logic, primary data source, optional enrichment resources, execution parameters, and export settings. SilkRoute interprets this descriptor to retrieve the initial record set and, when configured, expand it with structural, functional, pathway, chemical, or interaction information obtained from external repositories. Primary records and complementary data are preserved as separate source-aware outputs, while the original descriptor, execution metadata, generated-file manifest, and run summary document the acquisition context. The same workflow model supports protein-, compound-, and interaction-centered acquisition scenarios and produces traceable packages for downstream curation, analysis, archiving, and machine-learning applications.

The end-to-end case studies were complemented by an automated test suite covering descriptor validation, workflow routing, query interpretation, source-specific parsers, command-line behaviour, graphical descriptor handling, and core acquisition components.

At the time of evaluation, SilkRoute provided defined retrieval, enrichment, mapping, or interaction-oriented operations for 20 biological repositories (Table **2**). Nine of these source interfaces were exercised directly in the three case studies reported here, whereas the remaining interfaces extend the implemented acquisition coverage but were not used in the reported case-study workflows. The implemented resources cover protein sequences and annotations, predicted and experimental structures, functional classifications, pathways, molecular interactions, chemical entities, bioactivities, enzyme properties, biochemical reactions, kinetics, reference sequences, and proteomics records. A detailed summary of the implemented data services and their workflow roles is provided in Section S2 and Supplementary Table S2.

**Table 2:** Biological repositories implemented in SilkRoute, their workflow roles, and their use in the reported case studies.

| Source | Data type | Workflow role | Implemented use | Used in case studies |
| --- | --- | --- | --- | --- |
| UniProt [14] | Proteins | Primary retrieval | Protein queries and cross-references | Yes |
| AlphaFold DB [16] | Structures | Enrichment | Predicted protein structures | Yes |
| BioDBNet [49] | Identifiers | Mapping/enrichment | Cross-database identifier mapping | No |
| BioGRID [21] | Interactions | Interaction/enrichment | Protein interaction records | Yes |
| BRENDA [25] | Enzymes | Enrichment | Enzyme and biochemical metadata | No |
| ChEBI [24] | Chemical entities | Retrieval/enrichment | Chemical ontology records | No |
| ChEMBL [22] | Compound-s/bioactivity | Primary retrieval | Activity, assay, target, and compound records | Yes |
| Gene Ontology [17] | Functional terms | Query/enrichment support | Ontology-term metadata and biological-term resolution | Yes |
| InterPro [46] | Domains/families | Enrichment | Protein domain and family annotations | Yes |
| KEGG [50] | Pathways/reactions | Enrichment | Gene, pathway, compound, and reaction records | No |
| PANTHER [51] | Protein families | Enrichment | Family and functional classification | No |
| Pathway Commons [45] | Pathways/networks | Enrichment | Pathway and network records | Yes |
| PDB [15] | Structures | Enrichment | Experimental structure metadata and files | Yes |
| PRIDE [27] | Proteomics | Retrieval/enrichment | Proteomics project metadata | No |
| PubChem [23] | Compounds | Retrieval/enrichment | Compound properties and identifiers | No |
| Reactome [18] | Pathways | Enrichment | Pathway and reaction metadata | No |
| RefSeq [52] | Reference sequences | Retrieval/enrichment | Entrez-based sequence retrieval | No |
| Rhea [19] | Reactions | Enrichment | Biochemical reaction metadata | No |
| SABIO-RK [26] | Kinetics | Enrichment | Kinetic-law records | No |
| STRING [20] | Interactions | Interaction/enrichment | Protein interaction networks | Yes |

Repository support is workflow-specific. It indicates that selected endpoints, query operations, parsers, mapping procedures, or enrichment routines have been implemented for a defined acquisition role. It does not imply exhaustive coverage of every endpoint or data type exposed by the corresponding repository, nor that every implemented interface was exercised in the case studies presented in this article.

The interfaces exercised in the reported case studies span primary retrieval, cross-reference enrichment, ontology-based query resolution, structural and functional annotation, pathway acquisition, and interaction-oriented expansion. The remaining implemented interfaces provide additional acquisition options that were outside the scope of the three representative workflows.

The implemented resources can be combined according to their role in an acquisition task. UniProt [14], for example, can provide a primary protein set that is expanded with structural, functional, pathway, or interaction evidence, whereas ChEMBL [22] can provide activity-centered records that may be complemented with chemical information from PubChem [23] or ChEBI [24]. Interaction resources such as BioGRID [21] and STRING [20] produce relational outputs that remain distinct from the primary entity table. This role-based coverage allows SilkRoute to support different biomolecular contexts without requiring the participating repositories to share a common data structure.

The implemented functionality is accessible through a command-line interface, a Python API, and an optional graphical descriptor builder. The command-line interface was used to execute the complete case-study workflows from versioned YAML descriptors. The Python API returned acquired records and execution metadata in memory, supporting integration with scripts and notebooks. The graphical builder generated executable workflow-v1 descriptors from form-based inputs, after which the resulting files could be inspected, edited, versioned, and executed through the command-line interface. Representative examples of graphical descriptor generation, command-line execution, and programmatic execution are provided in Sections S3.2 and S5.1.2–S5.1.3 of the Supplementary Information.

These access paths support different intended uses. Command-line execution is suited to complete workflows that must be shared, archived, or rerun. Programmatic access supports acquisition embedded within custom analysis environments. The graphical builder assists users who prefer guided workflow specification while still producing an inspectable YAML artifact. The evaluation confirmed that each access path performed its intended function, although it was not designed as a formal usability or user-experience study.

### 3.2 Case-study evaluation across proteins, compounds, and interactions

Three representative workflows were selected to evaluate whether the SilkRoute acquisition model could support distinct biomolecular entities, query organizations, source relationships, and output structures. The protein-centered workflow combined primary retrieval with multi-source enrichment, the compound-centered workflow assembled a large bioactivity table from multiple query components, and the interaction-centered workflow expanded an initial protein set with relational evidence from external interaction resources. These scenarios assessed the framework across entity-level, measurement-level, and relationship-oriented acquisition tasks. Because the evaluation focused on software functionality, workflow execution, and acquisition traceability rather than on testing biological hypotheses, all reported quantities are descriptive outputs of the corresponding workflows and no inferential statistical analyses were performed. Detailed workflow configurations, execution examples, and representative outputs for the three case studies are provided in Section S5 of the Supplementary Information.

Table **3** summarizes the principal outputs obtained from the three case studies.

**Table 3:**
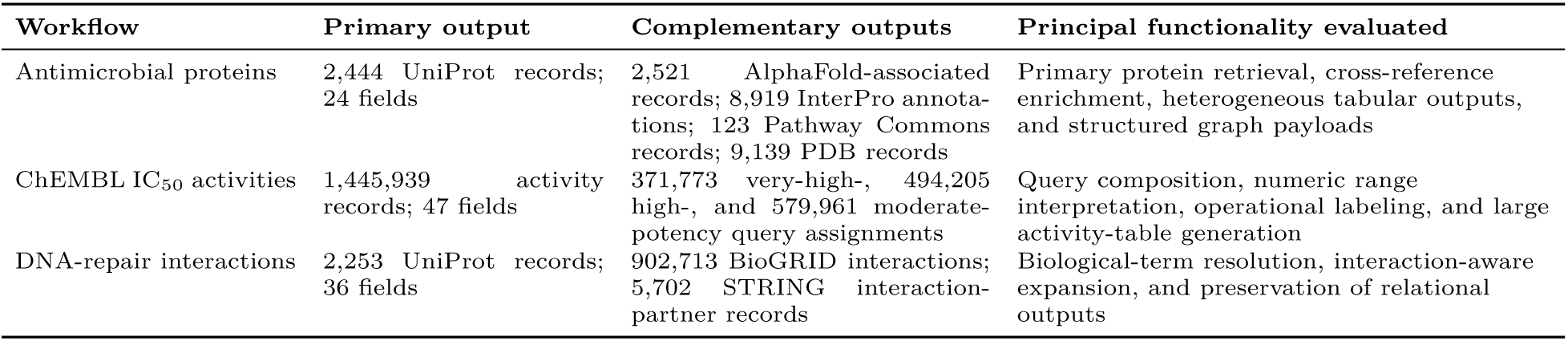
Summary of the representative workflows used to evaluate SilkRoute.

#### 3.2.1 Protein-centered acquisition and multi-source enrichment

The protein-centered workflow used UniProt as the primary source and retrieved reviewed protein records annotated with the term antimicrobial, with sequence lengths restricted to 100–500 amino acids. Available cross-references were subsequently used to obtain complementary information from AlphaFold DB, InterPro, Pathway Commons, and PDB. The complete workflow descriptor used for this case study is provided in Section S5.1.1 of the Supplementary Information.

The workflow retrieved 2,444 reviewed UniProt protein records, all of which were retained in the primary output. The exported table contained 24 fields, including accessions, protein names, organism information, sequences, sequence lengths, and cross-reference columns. The enrichment steps generated 2,521 AlphaFold-associated records, 8,919 InterPro annotation records, 123 Pathway Commons neighborhood records, and 9,139 PDB entry records. Each resource was exported separately, preserving the distinct structures and biological interpretations of the resulting annotations. Representative output files and their dimensions are summarized in Section S5.1.5 of the Supplementary Information.

The workflow also evaluated the handling of structured responses that could not be represented adequately in a conventional table. Pathway Commons neighborhood results were accompanied by compressed JSON files containing the raw graph payloads. The run produced 123 tabular neighborhood records and 123 non-empty graph files, with no missing or invalid payloads detected. Although the corresponding table contained zero-valued graph-count summaries, inspection of the associated files confirmed that the raw graph responses were present and structurally valid. The tabular records and compressed payloads should therefore be interpreted as complementary representations of the same enrichment operation. Further details on graph-payload storage and export are provided in Section S5.1.6 of the Supplementary Information.

The complete acquisition finished successfully in 14,095 seconds (approximately 3.9 hours) and generated the primary table, four enrichment outputs, structured graph payloads, execution metadata, and a run summary. This duration represents the observed wall-clock time of the reported run and should not be interpreted as a platform-independent performance benchmark. Network conditions, cache state, upstream service availability, repository response times, and changes in database content can all affect the duration of equivalent executions.

The protein case study demonstrates that SilkRoute can transform a compact biological query into a multi-source acquisition package linking primary protein records with predicted structures, experimental structures, functional annotations, and pathway-level information. More importantly, the resulting files retain the contribution of each repository, allowing downstream analyses to decide which forms of evidence should be merged, filtered, or retained independently.

#### 3.2.2 Compound bioactivity acquisition through query composition

The compound-centered workflow evaluated whether the acquisition model could support large measurement-oriented datasets that were not sequence-centered. ChEMBL was used as the primary source, and three labeled query components were defined for IC_50_ records: very_high_potency for 0 < IC50 < 10 nM, high_potency for 10 < IC50 < 100 nM, and moderate_potency for 100 < IC50 < 1000 nM. The intervals were defined as open ranges and therefore excluded the stated boundaries. The corresponding query-composition workflow and YAML descriptor are described in Sections S5.2.1–S5.2.2 of the Supplementary Information.

The workflow produced a single ChEMBL table containing 1,445,939 activity records and 47 fields. All retrieved records had standard_type = IC50 and standard_units = nM. The output contained 371,773 records assigned to the very_high_potency component, 494,205 assigned to high_potency, and 579,961 assigned to moderate_potency. The retrieved standard_value values ranged from 1.427 × 10−13 to 999.82 nM, with a mean value of 176.79 nM. No records had values exactly equal to 0, 10, 100, or 1000 nM, confirming that the open interval definitions were applied as specified. Representative output files and acquisition summaries are provided in Section S5.2.3 of the Supplementary Information.

The generated labels identify the query component through which each record was acquired. They should not be interpreted as validated pharmacological classes because the output remains a collection of source-derived ChEMBL activity records. Replicate aggregation, compound standardization, target-specific curation, conflict resolution, and final activity-label validation were not applied during acquisition.

Preserving the query-component assignment nevertheless provides a useful intermediate representation. It makes the numeric acquisition criteria explicit and allows subsequent curation procedures to determine whether the retrieved subsets should be retained, merged, relabeled, or restricted to particular assays, targets, compounds, or evidence conditions. The case study therefore shows that SilkRoute can support large activity-centered workflows while preserving the distinction between an operational acquisition label and a downstream biological interpretation.

#### 3.2.3 Interaction-aware protein dataset expansion

The interaction-centered workflow evaluated the acquisition of relational evidence. It began with a UniProt query for human proteins associated with DNA repair. The term DNA repair was resolved to the Gene Ontology identifier GO:0006281, resulting in the executable UniProt expression go:0006281 AND taxonomy_id:9606. The resulting protein set was subsequently expanded with interaction records from BioGRID and STRING.

The primary output contained 2,253 UniProt protein records and 36 fields. BioGRID produced 902,713 interaction records with 11 fields, while STRING generated 5,702 interaction-partner records with 18 fields. The interaction outputs were substantially larger than the initial protein table and were retained as independent files.

BioGRID requests used available gene symbols and organism identifiers from the primary records with taxId=9606. No evidence-type filter was configured, and self-interactions and interspecies interactions were not removed during acquisition. STRING requests used available STRING identifiers and otherwise fell back to gene symbols with species=9606. No workflow-level STRING confidence threshold or interaction-partner limit was applied. The resulting files should therefore be interpreted as broad acquisition outputs requiring downstream filtering, not as curated or confidence-filtered interaction networks. The complete interaction-aware workflow configuration, source-specific execution details, and representative outputs are provided in Sections S5.2.4–S5.2.6 of the Supplementary Information.

This case study illustrates why interaction evidence should remain structurally distinct from entity-level records. An interaction row describes a relationship between molecular entities and may contain repeated pairs, heterogeneous evidence types, source-specific identifiers, and database-specific confidence semantics. Collapsing these data into columns attached to individual protein records would remove relational information and obscure the operations required for deduplication, confidence filtering, or evidence integration. Preserving independent interaction outputs allows these decisions to be made according to the objectives of the downstream study.

### 3.3 Cross-workflow findings and generated acquisition packages

Across the three case studies, SilkRoute applied the same acquisition model to protein records, chemical activity measurements, and molecular interactions. The protein-centered workflow combined an initial record set with multiple enrichment resources; the ChEMBL workflow combined multiple numeric query components into a single activity table; and the interaction-centered workflow expanded an entity set into relational evidence whose size and structure differed substantially from the primary output.

Despite these differences, every case study produced a consistent set of acquisition artifacts comprising the original workflow descriptor, primary records, source-specific complementary outputs, execution metadata, and a compact run summary. The protein workflow additionally demonstrated the preservation of large structured payloads outside the principal tabular outputs. Figure **2** summarizes the resulting organization. Further details on the generated acquisition packages, their metadata structure, and representative workflow outputs are provided in Sections S4.2–S4.3 and S5 of the Supplementary Information.

**Figure 2:**
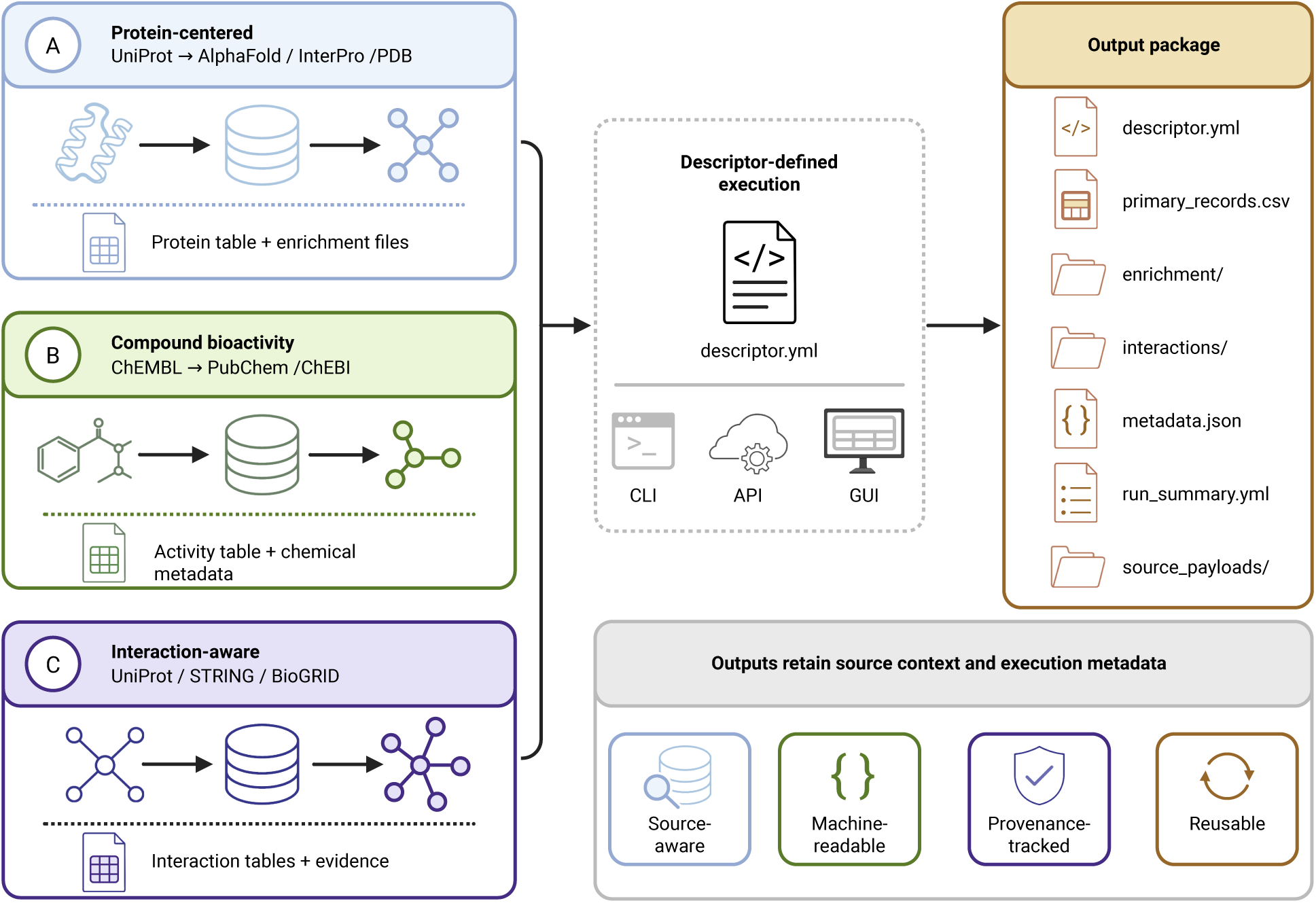
Representative SilkRoute workflows and generated output packages. Protein-centered, compound-bioactivity, and interaction-aware workflows are executed through a common descriptor-defined acquisition model. The resulting package preserves the workflow descriptor, primary records, enrichment files, interaction evidence when available, structured source payloads, execution metadata, and a run summary. The separation of these components keeps heterogeneous records source-aware and reusable for downstream curation, analysis, archiving, or machine-learning workflows. The source combinations shown illustrate supported workflow classes and do not reproduce every source or enrichment operation used in each reported case study.

Three findings were consistent across the evaluated workflows. First, descriptor-defined execution was applicable beyond protein-sequence retrieval and supported entity-level, measurement-level, and relational records. Second, source-aware output separation allowed structurally different data to be acquired without imposing premature schema unification. Third, the preserved descriptor and execution records connected each generated file with the query logic and source-specific operations that produced it.

The resulting artifact should therefore be interpreted as an acquisition package rather than as a fully curated or benchmark-ready dataset. Its principal value is that the transition from acquisition intent to retrieved records remains inspectable. Biological curation, filtering, harmonization, deduplication, labeling, and partitioning can subsequently be performed according to the scientific requirements of a particular study without first reconstructing the acquisition process from project-specific scripts or incomplete documentation.

This separation is especially relevant for collaborative or iterative dataset development. A versioned descriptor can be reviewed before execution, while the corresponding outputs and execution records can be archived as a release-specific acquisition package. When a repository is updated or acquisition criteria are revised, a new package can be generated without overwriting the configuration or outputs associated with the previous run. Further details on descriptor versioning, execution manifests, and reproducibility across repeated acquisitions are provided in Section S4.2 of the Supplementary Information.

### 3.4 Functional positioning relative to existing data-access approaches

Biopython and BioServices provide established foundations for accessing and processing biological information [34, 35], while repository-specific APIs provide direct access to individual databases. These approaches are complementary but operate at different levels. Biopython offers broad utilities for sequence processing, file parsing, and general bioinformatics analysis; BioServices provides programmatic clients for multiple biological web services; and direct APIs expose the endpoints and query capabilities of individual repositories.

SilkRoute is positioned at the level of dataset-oriented acquisition coordination. The comparison in Table **4** therefore focuses on capabilities provided directly as part of each approach, particularly those related to workflow specification, multi-source coordination, output organization, and execution reporting. In the table, “user-implemented” indicates that a capability can be developed using the corresponding library or service, but requires project-specific code or workflow logic and is not provided as a common dataset-level acquisition abstraction by default.

**Table 4:** Functional positioning of SilkRoute relative to established biological data-access approaches.

| Capability | SilkRoute | Biopython | BioServices | Repository-specific APIs |
| --- | --- | --- | --- | --- |
| Primary purpose | Multi-source dataset acquisition | General bioinformatics utilities | Access to biological web services | Direct repository access |
| Machine-readable dataset-level acquisition specification | Built-in `workflow-v1` descriptor | User-implemented | User-implemented | Source-specific or user-implemented |
| Explicit primary and enrichment roles | Built-in workflow concept | User-implemented | User-implemented | Requires cross-source coordination |
| Cross-source acquisition orchestration | Available for implemented workflows | User-implemented | User-implemented using service clients | User-implemented |
| Source-aware multi-file output package | Generated during complete execution | User-implemented | User-implemented | Limited to individual API responses |
| Workflow specification and normalised execution information preserved together | Generated by default | User-implemented | User-implemented | User-implemented |
| Metadata manifest and compact run summary | Generated by default | User-implemented | User-implemented | User-implemented |
| Command-line execution from a reusable YAML specification | Provided | No general dataset-acquisition model | No general dataset-acquisition model | Repository-dependent |
| Programmatic access | Python API | Python API | Python API | API- and language-dependent |
| Graphical workflow specification | Optional descriptor builder | Not provided as a common feature | Not provided as a common feature | Repository-dependent |
| General sequence-analysis and file-processing utilities | Outside core scope | Extensive | Limited | Outside API scope |
| Direct control over repository endpoints | Limited to implemented operations | Dependent on available clients | Dependent on available services | Provided directly |

The comparison is intended to clarify the level of abstraction provided by each approach and should not be interpreted as a ranking of their general capabilities. Biopython provides substantially broader functionality for sequence analysis, format handling, and general bioinformatics processing than SilkRoute. BioServices provides flexible access to a broad collection of biological web services that can be assembled into customized applications. Repository-specific APIs remain the most appropriate option when researchers require direct control over endpoints, parameters, or data types that are not implemented in SilkRoute.

These approaches can also be used to construct multi-source acquisition procedures. However, the coordination of source roles, query execution, cross-resource expansion, output organization, and provenance reporting generally remains part of the project-specific implementation. SilkRoute addresses this complementary problem by representing supported multi-source acquisition procedures as dataset-level workflows with explicit primary and enrichment roles, source-aware outputs, and associated execution records.

General-purpose workflow systems, including Galaxy, CWL-based platforms, and Nextflow, can provide execution portability, dependency management, scheduling, and provenance for arbitrary computational pipelines [36, 37, 38]. Data-recipe approaches can similarly support the preservation and reuse of data-processing procedures [39]. Their scope differs from that of SilkRoute because repository-specific acquisition tools, source relationships, output conventions, and biological query logic must still be defined or integrated within each workflow. SilkRoute instead provides a domain-specific acquisition model together with implemented biological source interfaces and conventions for organizing the resulting acquisition package. These approaches are complementary, and SilkRoute workflows could be incorporated into broader workflow-management systems when required.

The principal benefit of SilkRoute is therefore not broader repository coverage, unrestricted endpoint access, or universally faster retrieval. Its contribution is the reduction of project-specific coordination and reporting logic required to convert a biomolecular acquisition objective into an inspectable and reusable output package. This abstraction is particularly relevant when a workflow combines multiple repositories, must be reviewed by collaborators, is expected to be executed again for subsequent dataset releases, or provides the acquisition basis for a documented biological data resource.

A controlled speed comparison with Biopython, BioServices, or direct APIs was not considered informative for the present study. All approaches ultimately depend on external repository services, and wall-clock performance is strongly influenced by server availability, network conditions, rate limits, pagination behaviour, query size, caching, and the specific endpoints used. The reported protein-workflow runtime is therefore presented as an execution record and not as evidence of performance superiority.

### 3.5 Intended uses, limitations, and future development

SilkRoute is intended for researchers constructing biomolecular datasets from one or more external repositories. Representative uses include assembling the initial records for biological data resources, preparing source-aware inputs for curation pipelines, collecting structural or functional annotations, expanding molecular sets with interaction evidence, rebuilding acquisition packages after source updates, and documenting the upstream data collection that precedes machine-learning development.

Versioned descriptors can also function as shared acquisition protocols within collaborative projects. One researcher can define or review the acquisition criteria, another can execute the workflow, and downstream contributors can inspect the resulting files together with the recorded execution context. This arrangement reduces dependence on undocumented scripts, local command histories, or manually reconstructed download procedures. Further details on workflow descriptors, execution manifests, and their role in reproducible acquisition are provided in Section S4.2 of the Supplementary Information.

The benefits demonstrated by the case studies should be interpreted within the intended scope of the framework. SilkRoute does not perform expert biological curation, final label validation, compound standardization, advanced deduplication, leakage control, train–test partitioning, representation learning, or model benchmarking. A traceable acquisition package is therefore not equivalent to a biologically validated or benchmark-ready dataset. It provides the source records and execution context upon which those down-stream procedures can operate.

Dependence on external repositories remains the principal technical limitation. Databases may update annotations, change identifiers, alter endpoints or response schemas, impose rate limits, revise authentication requirements, or experience temporary interruptions [14, 29, 30, 53]. SilkRoute records how a workflow was configured and executed but cannot guarantee that a later execution will return identical biological content. Exact reconstruction of historical outputs additionally requires archived acquisition packages, stable database releases, or source snapshots.

Repository coverage is also intentionally selective. An implemented interface supports defined retrieval or enrichment operations and not necessarily every endpoint provided by the corresponding resource. Missing identifiers and incomplete cross-references can limit enrichment coverage, while large multi-source workflows can produce substantial runtimes and output volumes. Source-derived records may additionally require extensive harmonization or validation before they can be integrated into a downstream analytical dataset. Additional details on supported data services, implementation scope, and limitations associated with external repositories and incomplete metadata are provided in Sections S2 and S6.1 of the Supplementary Information.

Future development will focus on improving validation, reporting, and interoperability without extending the framework into downstream biological curation or benchmarking. Planned priorities include stricter descriptor validation, file checksums and release manifests, expanded validation of generated outputs, improved human-readable execution reports, and additional acquisition scenarios for peptides, enzyme properties, protein engineering, and compound-centered studies. Integration with broader dataset-documentation standards and general workflow-execution systems may further connect SilkRoute acquisition packages with downstream curation, release, and computational analysis pipelines. Additional planned directions are described in Section S6.2 of the Supplementary Information.

## 4 Conclusions

SilkRoute provides a descriptor-driven framework for reproducible biomolecular data acquisition from heterogeneous biological repositories. Across the protein-, compound-, and interaction-centered case studies, the same acquisition model successfully supported distinct biomolecular entity types, query organizations, enrichment strategies, and output structures. The evaluated workflows showed that primary records, complementary annotations, relational evidence, structured payloads, and execution metadata can be acquired and preserved within a common workflow specification without forcing heterogeneous source records into a single universal schema.

The main contribution of SilkRoute is the formalization of biomolecular data acquisition as an explicit and versionable component of dataset construction. Existing repositories, database clients, and programmatic APIs provide access to the underlying biological information, but the coordination of multi-source queries, enrichment steps, output organization, and execution reporting commonly remains embedded in project-specific scripts or manually documented procedures. SilkRoute addresses this workflow-level gap by connecting acquisition intent with source-aware outputs and provenance records that can be inspected, archived, reviewed, and reused.

This contribution is relevant because the reliability of a computational dataset depends not only on its final records, but also on the procedures used to select, retrieve, and expand them. Reproducible acquisition does not by itself guarantee biological validity, label quality, appropriate deduplication, or readiness for machine-learning evaluation. It does, however, provide a documented and auditable starting point for those downstream processes. By preserving both the acquisition specification and the resulting execution context, SilkRoute reduces the need to reconstruct dataset origins retrospectively and supports more transparent collaboration, resource development, dataset release, and computational analysis.

SilkRoute should therefore be viewed as an upstream acquisition and provenance layer that complements biological databases, curation procedures, and downstream analytical frameworks. Its descriptor-defined workflow model provides a practical foundation for constructing multi-source biomolecular datasets whose acquisition logic remains explicit, reusable, and traceable across studies and dataset versions.

## Declarations

### Availability and requirements

**Project name:** SilkRoute

**Project home page:** https://github.com/kren-ai-lab/silk-route

**Operating systems:** Linux and Windows

**Programming language:** Python

**Other requirements:** Python >=3.11,<3.15 and internet access for communication with external biological repositories. Authentication credentials or user identification may be required for selected data sources. Dependencies for the optional graphical descriptor builder are installed separately from the core package requirements.

**License:** MIT License

**Any restrictions to use by non-academics:** None

**Archived version** : SilkRoute v0.1.0, Zenodo, https://doi.org/10.5281/zenodo.21512367

## List of abbreviations

API: Application Programming Interface
CLI: Command-Line Interface
GUI: Graphical User Interface
YAML: YAML Ain’t Markup Language
JSON: JavaScript Object Notation
PDB: Protein Data Bank
GO: Gene Ontology
IC_50_: Half-maximal inhibitory concentration

## Ethics approval and consent to participate

Not applicable

## Consent for publication

Not applicable

## Availability of data and materials

The datasets and execution artifacts generated and/or analysed during the current study are publicly available in the SilkRoute v0.1.0 Zenodo archive [54] at https://doi.org/10.5281/zenodo.21512367. The archived materials include the representative workflow-v1 descriptors, primary and source-specific output files, structured payloads where applicable, execution metadata, run summaries, and the result artifacts supporting the protein-, compound-, and interaction-centered workflows reported in this article. All reported workflows and associated outputs were generated using SilkRoute v0.1.0.

The SilkRoute source code, documentation, installation instructions, and executable workflow examples are publicly available under the MIT License through the project repository at https://github.com/kren-ai-lab/silk-route. The package is also distributed through PyPI. Additional information on code availability and distribution is provided in Section S6.3 of the Supplementary Information.

## Authors’ contributions

DF, JG-V, FH-R, FC, and DM-O conceptualized the study. DF and DA-S implemented the software. DF, JG-V, DA-S, FH-R, and DM-O developed the methodology. FH-R and DM-O validated the framework and the representative workflows. JG-V, JM-F, MS-G, JS-Y, and XC conducted the investigation. MS-G designed and prepared the figures. MDD, RU-P, FH-R, and DM-O supervised the work and contributed research resources. FH-R and DM-O administered the project. MDD, RU-P, and DM-O contributed to funding acquisition. DF, JG-V, JM-F, MS-G, JS-Y, XC, FC, MDD, RU-P, FH-R, and DM-O contributed to writing, reviewing, and editing the manuscript. All authors read and approved the final manuscript.

## Acknowledgements

Not applicable.

## References

[1] Leman Binokay, Yavuz Oktay, and Gökhan Karakülah. The significance and evolution of biological databases in systems biology. In Systems Biology and In-Depth Applications for Unlocking Diseases, pages 137–148. Elsevier, 2025.

[2] Haoxing Luo, Yue Hu, Chaolin Song, Xinhui Li, Yuyin Ma, Yurong Qian, and Lei Deng. Me-pfp: An ensemble learning approach fusing multi-source features for protein function prediction. Journal of Chemical Information and Modeling, 2026.

[3] Misael Bessa Sales, Francisco Izaias da Silva Aires, Paulo Goncalves De Sousa, Calebe Da Rocha Silva, Emanuel Lino Rodrigues, Maria Cristiane Martins De Souza, and José CS dos Santos. Artificial intelligence in enzyme catalysis: Emerging trends and applications in biocatalyst engineering. The Canadian Journal of Chemical Engineering, 2026.

[4] Fang Sheng, Mohammad Noaeen, and Zahra Shakeri. Prodcarl: Reinforcement learning-aligned diffusion models for de novo antimicrobial peptide design. arXiv *preprint arXiv:2602.00157*, 2026.

[5] Edgar Lopez Lopez, Jean Paul Sánchez Castañeda, Massyel S Martinez-Cortés, Cesar de la Fuente-Nunez, and José Luis Medina-Franco. Exploring and expanding the chemical multiverse of peptides. Chemical Science, 2026.

[6] Areen Rasool, Jamshaid Ul Rahman, and Qasim Ali. Deephybridcpi: A hybrid deep learning framework for compound–protein interaction prediction. Journal of Molecular Graphics and Modelling, page 109303, 2026.

[7] Alec Lamens and Jürgen Bajorath. Explainable artificial intelligence for molecular design in pharmaceutical research. Chemical Science, 2026.

[8] David Medina-Ortiz, Sebastián Escobedo, Norma Murillo-Acevedo, Nicole Soto-García, Diego Fernández-Villegas, Diego Sandoval, and Anamaría Daza. Perspectives chapter: Data-centric strategies for machine learning-driven therapeutic peptide design – challenges and perspectives. In Data Quality Matters - Best Practices for Integrity and Assurance, chapter 23. IntechOpen, 2026.

[9] Vasileios Lapatas, Michalis Stefanidakis, Rafael C Jimenez, Allegra Via, and Maria Victoria Schneider. Data integration in biological research: an overview. Journal of Biological Research-Thessaloniki, 22(1):9, 2015.

[10] Fabio Herrera-Rocha, David Medina-Ortiz, Fabian Mauz, Juergen Pleiss, and Mehdi D Davari. Best practices for machine learning-assisted protein engineering. Journal of Chemical Information and Modeling, 65(23):12655–12667, 2025.

[11] Mark D Wilkinson, Michel Dumontier, IJsbrand Jan Aalbersberg, Gabrielle Appleton, Myles Axton, Arie Baak, Niklas Blomberg, Jan-Willem Boiten, Luiz Bonino da Silva Santos, Philip E Bourne, et al. The fair guiding principles for scientific data management and stewardship. Scientific data, 3(1):1–9, 2016.

[12] Fotis A Baltoumas, Sofia Zafeiropoulou, Evangelos Karatzas, Mikaela Koutrouli, Foteini Thanati, Kleanthi Voutsadaki, Maria Gkonta, Joana Hotova, Ioannis Kasionis, Pantelis Hatzis, et al. Biomolecule and bioentity interaction databases in systems biology: a comprehensive review. Biomolecules, 11(8):1245, 2021.

[13] Andrew D Rouillard, Gregory W Gundersen, Nicolas F Fernandez, Zichen Wang, Caroline D Monteiro, Michael G McDermott, and Avi Ma’ayan. The harmonizome: a collection of processed datasets gathered to serve and mine knowledge about genes and proteins. Database, 2016:baw100, 2016.

[14] The UniProt Consortium. Uniprot: the universal protein knowledgebase in 2025. Nucleic Acids Research, 53(D1):D609–D617, 2025.

[15] Helen M Berman and Stephen K Burley. Protein data bank (pdb): Fifty-three years young and having a transformative impact on science and society. Quarterly Reviews of Biophysics, 58:e9, 2025.

[16] Mihaly Varadi, Damian Bertoni, Paulyna Magana, Urmila Paramval, Ivanna Pidruchna, Malarvizhi Radhakrishnan, Maxim Tsenkov, Sreenath Nair, Milot Mirdita, Jingi Yeo, et al. Alphafold protein structure database in 2024: providing structure coverage for over 214 million protein sequences. Nucleic Acids Research, 52(D1):D368–D375, 2024.

[17] Gene Ontology Consortium. The gene ontology knowledgebase in 2026. Nucleic Acids Research, 54(D1):D1779–D1792, 2026.

[18] Marija Milacic, Deidre Beavers, Patrick Conley, Chuqiao Gong, Marc Gillespie, Johannes Griss, Robin Haw, Bijay Jassal, Lisa Matthews, Bruce May, et al. The reactome pathway knowledgebase 2024. Nucleic Acids Research, 52(D1):D672–D678, 2024.

[19] Parit Bansal, Anne Morgat, Kristian B Axelsen, Venkatesh Muthukrishnan, Elisabeth Coudert, Lucila Aimo, Nevila Hyka-Nouspikel, Elisabeth Gasteiger, Arnaud Kerhornou, Teresa Batista Neto, et al. Rhea, the reaction knowledgebase in 2022. Nucleic Acids Research, 50(D1):D693–D700, 2022.

[20] Damian Szklarczyk, Katerina Nastou, Mikaela Koutrouli, Rebecca Kirsch, Farrokh Mehryary, Radja Hachilif, Dewei Hu, Matteo E Peluso, Qingyao Huang, Tao Fang, et al. The string database in 2025: protein networks with directionality of regulation. Nucleic Acids Research, 53(D1):D730–D737, 2025.

[21] Rose Oughtred, Jennifer Rust, Christie Chang, Bobby-Joe Breitkreutz, Chris Stark, Andrew Willems, Lorrie Boucher, Genie Leung, Nadine Kolas, Frederick Zhang, et al. The biogrid database: A comprehensive biomedical resource of curated protein, genetic, and chemical interactions. Protein Science, 30(1):187–200, 2021.

[22] Barbara Zdrazil, Eloy Felix, Fiona Hunter, Emma J Manners, James Blackshaw, Sybilla Corbett, Marleen De Veij, Harris Ioannidis, David Mendez Lopez, Juan F Mosquera, et al. The chembl database in 2023: a drug discovery platform spanning multiple bioactivity data types and time periods. Nucleic Acids Research, 52(D1):D1180–D1192, 2024.

[23] Sunghwan Kim, Jie Chen, Tiejun Cheng, Asta Gindulyte, Jia He, Siqian He, Qingliang Li, Benjamin A Shoemaker, Paul A Thiessen, Bo Yu, et al. Pubchem 2025 update. Nucleic Acids Research, 53(D1):D1516–D1525, 2025.

[24] Adnan Malik, Muhammad Arsalan, Carlos Moreno, Juan Mosquera, Eloy Félix, Tevfik Kizilören, Venkatesh Muthukrishnan, Barbara Zdrazil, Andrew R Leach, and Noel M O’Boyle. Chebi: reengineered for a sustainable future. Nucleic Acids Research, 54(D1):D1768–D1778, 2026.

[25] Julia Hauenstein, Lisa Jeske, Antje Jäde, Mathias Krull, Katrin Dümmer, Julia Koblitz, Anja Tietz, Dieter Jahn, Lorenz Christian Reimer, and Boyke Bunk. Brenda in 2026: a global core biodata resource for functional enzyme and metabolic data within the dsmz digital diversity. Nucleic Acids Research, 54(D1):D527–D534, 2026.

[26] Ulrike Wittig, Maja Rey, Andreas Weidemann, Renate Kania, and Wolfgang Müller. Sabio-rk: an updated resource for manually curated biochemical reaction kinetics. Nucleic Acids Research, 46(D1):D656–D660, 2018.

[27] Yasset Perez-Riverol, Chakradhar Bandla, Deepti J Kundu, Selvakumar Kamatchinathan, Jingwen Bai, Suresh Hewapathirana, Nithu Sara John, Ananth Prakash, Mathias Walzer, Shengbo Wang, et al. The pride database at 20 years: 2025 update. Nucleic Acids Research, 53(D1):D543–D553, 2025.

[28] Mark Ziemann, Pierre Poulain, and Anusuiya Bora. The five pillars of computational reproducibility: bioinformatics and beyond. Briefings in Bioinformatics, 24(6):bbad375, 2023.

[29] Eric W Sayers, Jeff Beck, Evan E Bolton, J Rodney Brister, Jessica Chan, Donald C Comeau, Ryan Connor, Michael DiCuccio, Catherine M Farrell, Michael Feldgarden, et al. Database resources of the national center for biotechnology information. Nucleic Acids Research, 52(D1):D33–D43, 2024.

[30] Mihai Pop, Teresa K Attwood, Judith A Blake, Philip E Bourne, Ana Conesa, Terry Gaasterland, Lawrence Hunter, Carl Kingsford, Oliver Kohlbacher, Thomas Lengauer, et al. Biological databases in the age of generative artificial intelligence. Bioinformatics Advances, 5(1):vbaf044, 2025.

[31] Judith Bernett, David B Blumenthal, Dominik G Grimm, Florian Haselbeck, Roman Joeres, Olga V Kalinina, and Markus List. Guiding questions to avoid data leakage in biological machine learning applications. Nature Methods, 21(8):1444–1453, 2024.

[32] Benjamin J Heil, Michael M Hoffman, Florian Markowetz, Su-In Lee, Casey S Greene, and Stephanie C Hicks. Reproducibility standards for machine learning in the life sciences. Nature methods, 18(10):1132–1135, 2021.

[33] João Felipe Pimentel, Leonardo Murta, Vanessa Braganholo, and Juliana Freire. A large-scale study about quality and reproducibility of jupyter notebooks. In 2019 IEEE/ACM 16th International Conference on Mining Software Repositories (MSR), pages 507–517. IEEE, 2019.

[34] Peter JA Cock, Tiago Antao, Jeffrey T Chang, Brad A Chapman, Cymon J Cox, Andrew Dalke, Iddo Friedberg, Thomas Hamelryck, Frank Kauff, Bartek Wilczynski, et al. Biopython: freely available python tools for computational molecular biology and bioinformatics. Bioinformatics, 25(11):1422, 2009.

[35] Thomas Cokelaer, Dennis Pultz, Lea M Harder, Jordi Serra-Musach, and Julio Saez-Rodriguez. Bioservices: a common python package to access biological web services programmatically. Bioinformatics, 29(24):3241–3242, 2013.

[36] The galaxy platform for accessible, reproducible, and collaborative data analyses: 2024 update. Nucleic acids research, 52(W1):W83–W94, 2024.

[37] Michael R Crusoe, Sanne Abeln, Alexandru Iosup, Peter Amstutz, John Chilton, Nebojša Tijanić, Hervé Ménager, Stian Soiland-Reyes, Bogdan Gavrilović, Carole Goble, et al. Methods included: standardizing computational reuse and portability with the common workflow language. Communications of the ACM, 65(6):54–63, 2022.

[38] Paolo Di Tommaso, Maria Chatzou, Evan W Floden, Pablo Prieto Barja, Emilio Palumbo, and Cedric Notredame. Nextflow enables reproducible computational workflows. Nature biotechnology, 35(4):316–319, 2017.

[39] Qian Liu, Qiang Hu, Song Liu, Alan Hutson, and Martin Morgan. Reusedata: an r/bioconductor tool for reusable and reproducible genomic data management. BMC bioinformatics, 25(1):8, 2024.

[40] Sean R Wilkinson, Meznah Aloqalaa, Khalid Belhajjame, Michael R Crusoe, Bruno de Paula Kinoshita, Luiz Gadelha, Daniel Garijo, Ove Johan Ragnar Gustafsson, Nick Juty, Sehrish Kanwal, et al. Applying the fair principles to computational workflows. Scientific data, 12(1):328, 2025.

[41] Samantha Petti and Sean R Eddy. Constructing benchmark test sets for biological sequence analysis using independent set algorithms. PLOS Computational Biology, 18(3):e1009492, 2022.

[42] Lei Wang, Xudong Li, Han Zhang, Jinyi Wang, Dingkang Jiang, Zhidong Xue, and Yan Wang. A comprehensive review of protein language models. arXiv preprint arXiv:2502.06881, 2025.

[43] Marine Djaffardjy, George Marchment, Clémence Sebe, Raphael Blanchet, Khalid Belhajjame, Alban Gaignard, Frédéric Lemoine, and Sarah Cohen-Boulakia. Developing and reusing bioinformatics data analysis pipelines using scientific workflow systems. Computational and Structural Biotechnology Journal, 21:2075–2085, 2023.

[44] Shadab Ahmad, Leonardo Jose da Costa Gonzales, Emily H Bowler-Barnett, Daniel L Rice, Minjoon Kim, Supun Wijerathne, Aurélien Luciani, Swaathi Kandasaamy, Jie Luo, Xavier Watkins, et al. The uniprot website api: facilitating programmatic access to protein knowledge. Nucleic Acids Research, 53(W1):W547–W553, 2025.

[45] Igor Rodchenkov, Ozgun Babur, Augustin Luna, Bulent Arman Aksoy, Jeffrey V Wong, Dylan Fong, Max Franz, Metin Can Siper, Manfred Cheung, Michael Wrana, et al. Pathway commons 2019 update: integration, analysis and exploration of pathway data. Nucleic Acids Research, 48(D1):D489–D497, 2020.

[46] Matthias Blum, Antonina Andreeva, Laise Cavalcanti Florentino, Sara Rocio Chuguransky, Tiago Grego, Emma Hobbs, Beatriz Lazaro Pinto, Ailsa Orr, Typhaine Paysan-Lafosse, Irina Ponamareva, et al. Interpro: the protein sequence classification resource in 2025. Nucleic Acids Research, 53(D1):D444–D456, 2025.

[47] Kirill Simonov. Pyyaml: Yaml parser and emitter for python. GitHub Repository, 2024.

[48] Ritchie Vink, Alexander Beedie, Orson Peters, Gijs Burghoorn, Stijn de Gooijer, Marco Edward Gorelli, et al. pola-rs/polars: Python polars 1.43.0, 2026.

[49] Uma Mudunuri, Anney Che, Ming Yi, and Robert M Stephens. biodbnet: the biological database network. Bioinformatics, 25(4):555–556, 2009.

[50] Minoru Kanehisa, Miho Furumichi, Yoko Sato, Yuriko Matsuura, and Mari Ishiguro-Watanabe. Kegg: biological systems database as a model of the real world. Nucleic Acids Research, 53(D1):D672–D677, 2025.

[51] Paul D Thomas, Dustin Ebert, Anushya Muruganujan, Tremayne Mushayahama, Laurent-Philippe Albou, and Huaiyu Mi. Panther: Making genome-scale phylogenetics accessible to all. Protein Science, 31(1):8–22, 2022.

[52] Tamara Goldfarb, Vamsi K Kodali, Shashikant Pujar, Vyacheslav Brover, Barbara Robbertse, Catherine M Farrell, Dong-Ha Oh, Alexander Astashyn, Olga Ermolaeva, Diana Haddad, et al. Ncbi refseq: reference sequence standards through 25 years of curation and annotation. Nucleic Acids Research, 53(D1):D243–D257, 2025.

[53] Mohamed Helmy, Alexander Crits-Christoph, and Gary D Bader. Ten simple rules for developing public biological databases. PLoS computational Biology, 12(11):e1005128, 2016.

[54] Diego Fernández, Julián A. García-Vinuesa, Diego Alvarez-Saravia, José L. Medina-Franco, Julieta H. Sepulveda-Yanez, Xavier Cadet, Frederic Cadet, Mehdi D. Davari, Roberto Uribe-Paredes, Fabio Herrera-Rocha, and David Medina-Ortiz. SilkRoute Case Studies: Workflow configurations, biomolecular datasets, and provenance metadata, 2026. Dataset.

