## Supplementary Information for "SilkRoute: A Descriptor-Driven Framework for Reproducible Multi-Source Biomolecular Data Acquisition"

[1]Departamento de Ingeniería en Computación, Universidad de Magallanes, Avenida Bulnes 01855, Punta Arenas, Chile.
[2]DIFACQUIM Research Group, Department of Pharmacy, School of Chemistry, Universidad Nacional Autónoma de México, Avenida Universidad 3000, 04510, Mexico City, Mexico
[3]Facultad de Ciencias de la Salud, Universidad de Magallanes, Avenida Bulnes 01855, Punta Arenas, Chile.
[4]Centro Asistencial de Docencia e Investigación, CADI, Universidad de Magallanes, Av. Los Flamencos 01364, 6210005, Punta Arenas, Chile.
[5]Dartmouth College, Hanover, NH, 03755, USA
[6]PEACCEL, AI for Biologics, Paris, France
[7]Leibniz-Institute of Plant Biochemistry, Department of Bioorganic Chemistry, Weinberg 3, D-06120 Halle, Germany
† These authors contributed equally
†† Work done whilst at Imperial College London, UK

#### S1 Software Architecture and Implementation Details

SilkRoute implements biomolecular data acquisition workflows through a descriptor-driven architecture. A user provides a YAML workflow descriptor, which is validated against the `workflow-v1` schema and converted into normalized workflow values before execution. These normalized values define the dataset modality, execution mode, executable query, enrichment options, harmonization fields, and export behavior.

The workflow execution layer then routes the request to the appropriate source-specific interfaces. Each interface encapsulates the details required to communicate with one biological resource, including endpoint configuration, request parameters, identifier handling, response parsing, and caching when available. This design keeps database- specific logic separate from the workflow layer, while allowing the workflow layer to coordinate query execution, enrichment, metadata capture, and export.

The end-to-end execution flow can be summarized as follows:

```
YAML descriptor → validation → normalized workflow values → source-specific interface → parser
→ enrichment → export and reporting → metadata.json/run_summary.yml
```

Primary records are first retrieved from the source defined by the workflow mode and query. When enrichment is enabled, cross-reference fields are used to request additional source specific records, such as AlphaFold metadata, InterPro entries, PathwayCommons neighborhood graphs, PDB entries, ChEMBL activities, or interaction records, depending on the workflow configuration. These enrichment outputs are exported as independent tables rather than being forced into a single denormalized table. This avoids hiding the heterogeneous structure of the original sources and preserves the interpretation of each output file.

When supported by the source-specific output, enrichment tables may include source-context columns that link enriched records back to the primary query results and to the external source from which they were obtained. These source-context columns are produced by the corresponding source-specific interfaces. The descriptor-level `harmonization` section does not independently create cross-source links or normalize heterogeneous output schemas.

The export layer writes the resulting tables together with machine-readable execution metadata. The original descriptor records the intended workflow configuration, whereas `metadata.json` and `run_summary.yml` describe the observed execution, including normalized parameters, output files, record counts, execution status, and source-specific metadata when available. This separation makes the workflow easier to inspect, repeat, and audit.

SilkRoute is implemented in Python. Tabular outputs are represented with `polars`, YAML descriptors are parsed with `PyYAML`, HTTP-based services are accessed through source-specific request logic, and the command-line interface provides descriptor-level workflow execution. The implementation is organized so that database-specific request and parsing logic remains inside dedicated interfaces, while workflow validation, execution, reporting, and export are coordinated by the higher-level workflow layer.

The following section describes the practical installation requirements, external service configuration, and supported data services used by this architecture.

#### S2 Installation, External Configuration, and Supported Data Services

SilkRoute can be installed in a standard Python environment. For most users, the recommended installation method is through PyPI:

```
pip install silkroute
```

Alternatively, the package can be installed from a local source checkout, which is useful for development, testing, or working with the current repository version. A typical source-based installation consists of creating an isolated environment and installing the package in editable mode:

```
conda create -n silkroute python=3.13
conda activate silkroute
pip install -e .
```

The analyses reported in this study were performed using SilkRoute v0.1.0 in isolated Python environments.

After installation, the command-line interface can be inspected with:

```
silkroute --help
```

Most source-specific configuration files required by SilkRoute are distributed with the package and are loaded automatically at runtime. Therefore, users do not need to manually copy field maps or endpoint configuration files before running the standard workflows. The main user-facing configuration concerns credentials for external services that require authentication or user identification.

**Supplementary Table S1**: External service configuration used by selected SilkRoute interfaces.

| Service | Requirement | Environment variable |
|---|---|---|
| BioGRID | API access key required for BioGRID interaction queries. | `SILKROUTE_BIOGRID_API_KEY` |
| BRENDA | Registered account credentials required for selected SOAP-based enzyme queries. | `SILKROUTE_BRENDA_EMAIL` `SILKROUTE_BRENDA_PASSWORD` |
| RefSeq / Entrez | User email recommended or required for NCBI Entrez-based requests. | `SILKROUTE_REFSEQ_EMAIL` |

Credentials can be provided as environment variables or through a local `.env` file. Such files should not be committed to version control. Workflows that rely only on open HTTP endpoints, such as the protein example based on UniProt, AlphaFold, InterPro, PathwayCommons, and PDB metadata retrieval, do not require BioGRID or BRENDA credentials. In contrast, interaction-focused workflows that query BioGRID require a valid BioGRID access key.

This distinction is important for reproducibility: the YAML descriptor defines the intended workflow, but successful execution also depends on the availability of the external services used by that workflow and, when applicable, on the presence of valid credentials.

SilkRoute provides implemented interfaces and configured retrieval routines for multiple biological data resources. In this context, *supported* means that the software implements one or more source-specific methods, endpoints, query builders, or parsers for the named resource.

The supported sources cover protein sequence retrieval, structure-related metadata, functional annotation, interaction evidence, pathway context, compound bioactivity, enzyme metadata, identifier mapping, and reference/proteomics records. Table **S2** summarizes the role of each source within SilkRoute workflows.

**Supplementary Table S2**: Biological data services with implemented SilkRoute support.

| Category | Source | Contribution to a workflow | Implementation scope |
|---|---|---|---|
| Primary protein records | UniProt (Release June 09, 2026) [1] | Protein sequences, accessions, annotations, taxonomy, features, and cross-references used as the main entry point for protein workflows. | Primary protein query source and cross-reference provider. |
| Reference sequences | RefSeq (Release May 19, 2026) [2] | Reference sequence records retrieved through NCBI services. | Entrez-based access; user email configuration may be required or recommended. |
| Protein structure | AlphaFold DB (Release 2025) [3] | Predicted protein-model metadata and optional model-file acquisition linked to UniProt accessions. | Selected prediction and download endpoints. |
| Protein structure | PDB (Date July 08, 2026) [4] | Experimental structure metadata and optional structure-file acquisition from RCSB PDB. | Selected entry metadata and download operations. |
| Protein annotation | InterPro (Release June 10, 2026) [5] | Protein families, domains, signatures, and associated functional annotations. | UniProt-linked entry retrieval and annotation parsing. |
| Protein annotation | PANTHER (Release June 20, 2024) [6] | Protein family, orthology, functional, and evolutionary classifications. | Selected classification endpoints. |
| Protein annotation | Gene Ontology (Release June 19, 2026) [7] | Ontology terms and gene-product annotations used to describe molecular function, biological process, and cellular component information. | Selected ontology and annotation endpoints. |
| Pathway and network context | Pathway Commons (Release july 16, 2024) [8] | Integrated pathway neighborhood and graph context associated with protein queries. | Selected fetch, top-pathway, and neighborhood operations; graph payloads may be exported externally. |
| Pathway and network context | Reactome (Release June 23, 2026) [9] | Pathway, reaction, event, entity, disease, and diagram metadata. | Selected Content Service and exporter endpoints. |
| Pathway and network context | KEGG (Release July 01, 2026) [10] | Genes, compounds, pathways, reactions, diseases, drugs, modules, and orthology records. | Selected KEGG REST operations. |
| Protein interactions | BioGRID (Release March 01, 2026) [11] | Genetic and physical interaction records associated with queried proteins or identifiers. | Credentialed API; selected interaction retrieval. |
| Protein interactions | STRING (Release July 26, 2023) [12] | Functional protein association records, interaction partners, scores, and source-specific evidence fields. | Selected interaction partner retrieval; STRING confidence scores retain source semantics. |
| Chemical entities | ChEBI (Realease July 07, 2026) [13] | Chemical entities and ontology relationships. | Selected entity searches and configured fields. |
| Chemical entities and assays | PubChem (Realease July 02, 2026) [14] | Compound, substance, assay, protein, gene, pathway, and taxonomy records. | Selected PUG-REST and PUG-View operations. |
| Compound bioactivity | ChEMBL (Realease May 01, 2026) [15] | Molecules, targets, assays, and bioactivity records, including activity-based workflows. | Endpoint-specific retrieval, filtering, pagination, and activity parsing. |
| Enzyme information | BRENDA (Realease March 04, 2026) [16] | Enzyme, kinetic, substrate, cofactor, inhibitor, pH, and temperature metadata. | Selected credentialed SOAP methods. |
| Enzyme kinetics | SABIO-RK (Realease May 19, 2026) [17] | Enzyme kinetic laws and experimental condition metadata. | Selected kinetic-law retrieval. |
| Biochemical reactions | Rhea (Realease June 10, 2026) [18] | Curated biochemical reaction records. | Reaction-centric configured fields and endpoints. |

*Continued on next page*

| Category | Source | Contribution to a workflow | Implementation scope |
|---|---|---|---|
| Identifier mapping | BioDBNet (Realease July 05, 2026) [19] | Identifier conversion between supported biological identifier systems. | Mapping utility rather than a primary annotation table. |
| Proteomics metadata | PRIDE (Realease July 08, 2026) [20] | Proteomics project and dataset metadata. | Project-level acquisition, not raw spectrum processing. |

Returned coverage depends on the query, identifier availability, external database state, authentication, API limits, and the fields populated by each upstream source at execution time. Therefore, a missing enrichment row should not be interpreted as direct biological evidence of absence. It may indicate that a cross-reference was not present, that the queried source did not expose a compatible record, or that the external request did not return data for that identifier.

The examples in this supplementary material use only a subset of these sources. The protein workflow uses UniProt as the primary query source and enriches the retrieved proteins with AlphaFold, InterPro, Pathway Commons, and PDB records. The compound workflow uses ChEMBL activity records, while the interaction-aware workflow combines UniProt-derived proteins with interaction evidence from BioGRID and STRING.

After defining the installation requirements and available data services, the next section describes how these choices are encoded in reproducible `workflow-v1` descriptors.

#### S3 Workflow Descriptor Schema

SilkRoute workflows are defined through YAML descriptors that follow the `workflow-v1` schema. The descriptor is the central reproducibility artifact of a workflow: it records the intended dataset, execution mode, executable query, enrichment configuration, harmonization fields, and export behavior before the workflow is run.

A descriptor acts as a compact protocol for dataset acquisition, allowing the same workflow definition to be reviewed, versioned, shared, and re-executed. During execution, SilkRoute validates the descriptor, normalizes user-facing values into internal workflow parameters, executes the requested source-specific operations, and writes both tabular outputs and machine-readable metadata.

Table **S3** summarizes the major sections of a `workflow-v1` descriptor.

**Supplementary Table S3**: Major sections of a SilkRoute `workflow-v1` descriptor.

| Section | Purpose |
|---|---|
| `schema_version` | Declares the descriptor schema used by the workflow. For the workflows reported here, this value is `workflow-v1`. |
| `dataset` | Stores dataset-level metadata, including the dataset name, description, biological modality, execution mode, and interaction type when applicable. |
| `query` | Defines the executable query and the fields requested from the primary source. It also specifies cross-reference fields used for enrichment. |
| `execution` | Controls runtime behavior, including enrichment, retry settings, worker count, debugging, pagination, timeouts, and optional structure-file downloads. |
| `harmonization` | Declares optional semantic column roles and limited export or reporting behavior. The `id_column` field can add deterministic row identifiers to exported tabular files, while `sequence_column` can support unique-sequence reporting. Other fields are preserved as descriptor metadata and do not transform source-specific schemas. |
| `export` | Defines the output directory, export format, metadata files, run summary files, and optional handling of large graph payloads. |

The following minimal descriptor illustrates the general structure of a `workflow-v1` file. Placeholder values are shown between angle brackets and should be replaced by workflow-specific values.

```
schema_version: workflow-v1
```

```yaml
dataset:
  name: <dataset_name>
  description: <short_dataset_description>
  modality: <protein|compound|interaction>
  mode: <query_first|query_composition>

query:
  value: <executable_query>
  fields:
    - <primary_field_1>
    - <primary_field_2>
  crossref_fields:
    - <enrichment_source_1>
    - <enrichment_source_2>
  include_isoform: false

execution:
  enrich: true
  max_workers: 5
  total_retries: 3
  debug: false

harmonization:
  metadata_fields:
    - <identifier_field>
    - <context_field>

export:
  output_dir: <output_directory>
  format: csv
  include_metadata: true
  include_summary: true
  manifest_file: metadata.json
  summary_file: run_summary.yml
```

Listing 1: Minimal SilkRoute workflow descriptor with placeholders.

The `query.value` field is the executable query used by the workflow. Other query-related fields define what should be requested or enriched, but the workflow is ultimately driven by the executable expression stored in `query.value`. For this reason, the descriptor should be considered the authoritative specification of the workflow.

The `execution` section controls how the workflow is run, but it does not change the biological meaning of the query. For example, increasing `max_workers` may change runtime behavior, while enabling `enrich` determines whether cross-reference-based enrichment outputs are requested. Similarly, the `export` section determines how outputs are written without changing the upstream query itself.

##### S3.1 Query Modes and Executable Query Fields

SilkRoute workflows separate the execution mode from the query expression itself. The execution mode defines how the workflow interprets the query and organizes the retrieved records, whereas `query.value` stores the executable expression passed to the corresponding query builder or source-specific interface.

The workflows reported in this supplementary material use two main query modes: `query_first` and `query_composition`. Their roles are summarized in Table **S4**.

**Supplementary Table S4**: Query modes used by SilkRoute workflow descriptors.

| Mode | Description |
|---|---|
| `query_first` | Executes a single primary query and uses the resulting records as the starting point for optional enrichment. This mode is used for the protein workflow example, where UniProt records are retrieved first and then enriched with AlphaFold, InterPro, PathwayCommons, and PDB records. |
| `query_composition` | Executes or interprets multiple query components that define labeled subsets or acquisition groups. This mode is used for activity-based ChEMBL workflows, where different activity ranges can be associated with different labels. |

The `query.value` field is the executable query field. In a `query_first` workflow, it defines the primary search expression that will be interpreted and submitted to the corresponding source-specific interface. For example, the protein workflow uses the following UniProt-oriented query:

```
antimicrobial AND reviewed:true AND length:100-500
```

SilkRoute includes a query interpretation layer that translates selected user-facing expressions into source-specific query syntax. This layer is not intended to be a general biological reasoning system or a complete ontology lookup service. Instead, it applies explicit transformations implemented for supported fields, such as range rewriting, alias handling, and source-specific query normalization.

In the antimicrobial protein example, the user-facing length filter:

```
length:100-500
```

is converted before submission to UniProt into the range syntax expected by the UniProt REST query language:

```
length:[100 TO 500]
```

Thus, the complete query stored in the YAML descriptor:

```
antimicrobial AND reviewed:true AND length:100-500
```

is normalized at execution time into a UniProt-compatible expression:

```
antimicrobial AND reviewed:true AND length:[100 TO 500]
```

In this case, `antimicrobial` is preserved as a free-text query term, `reviewed:true` is already compatible with the UniProt query syntax, and `length:100-500` is rewritten as a UniProt range expression. The original descriptor therefore preserves a compact and readable query, while the execution metadata records the normalized query parameters observed during runtime.

The same interpretation principle is used in other source-specific workflows, although the target syntax depends on the database being queried. For example, in ChEMBL activity workflows, a compact IC50 expression such as:

```
ic50:100-1000
```

can be interpreted as an activity-oriented filter over IC50 records:

```
standard_type=IC50 AND standard_value>100 AND standard_value<1000
```

Similarly, threshold-style IC50 expressions can be rewritten into explicit ChEMBL activity filters. For example:

```
ic50:<10
```

corresponds to an IC50 activity query constrained by:

```
standard_type=IC50 AND standard_value<10
```

In `query_composition` workflows, `query.value` may contain multiple query components, each representing a subset or label. For example, a ChEMBL activity workflow can define several IC50 ranges and associate each range with a label used in the exported activity table:

```
ic50:0-10=very_high_potency,
ic50:10-100=high_potency,
ic50:100-1000=moderate_potency
```

This allows a single descriptor to document both the acquisition logic and the label assignment strategy used during retrieval. These labels should be interpreted as descriptor-defined acquisition labels rather than independently curated biological classes unless additional downstream validation is performed.

The `query.fields` entry controls which fields are requested from the primary source when the interface supports field selection. In protein workflows, these fields may include accessions, protein names, organism metadata, sequences, taxonomy identifiers, and cross-reference fields. The `query.crossref_fields` entry controls which additional enrichment sources should be queried after the primary records have been retrieved.

For example, the following query configuration requests primary UniProt fields and enables enrichment through AlphaFold, PDB, PathwayCommons, and InterPro cross-references:

```
query:
  value: antimicrobial AND reviewed:true AND length:100-500
  fields:
    - accession
    - protein_name
    - organism_name
    - sequence
    - organism_id
  crossref_fields:
    - alphafold
    - pdb
    - pathwaycommons_neighborhood
    - interpro
  include_isoform: false
```

The `fields` and `crossref_fields` entries therefore have different roles. The first controls the primary record structure requested from the initial source, whereas the second controls which enrichment outputs are generated from the retrieved records. This distinction is important because enrichment outputs are exported as source-specific tables rather than being merged into a single universal table.

For interaction-aware workflows, the descriptor may also include an `interaction_type` field in the `dataset` section. This field documents whether the workflow is oriented toward protein–protein or protein–ligand interaction evidence. The interaction type helps interpret the workflow and route execution, but the query itself remains defined by `query.value`.

The interpreter provides a controlled bridge between compact descriptor-level queries and the syntax expected by supported external sources. Unsupported terms are not semantically inferred by SilkRoute and must either be valid for the target source or be handled by the corresponding source-specific interface.

##### S3.2 Graphical User Interface for YAML Descriptor Generation

SilkRoute also provides a graphical user interface for generating `workflow-v1` YAML descriptors. The GUI is intended to reduce the manual effort required to write workflow configuration files and to make the descriptor structure more accessible to users who are less familiar with YAML syntax.

The graphical interface exposes the main descriptor sections, including `dataset`, `query`, `execution`, `harmonization`, and `export`. User-facing labels are mapped to the internal values expected by the workflow schema. For example, a visible option such as *Query First* is exported as `query_first`, and selected enrichment resources are written under `query.crossref_fields`. The generated file can then be inspected, versioned, edited manually, or executed through the command-line workflow runner.

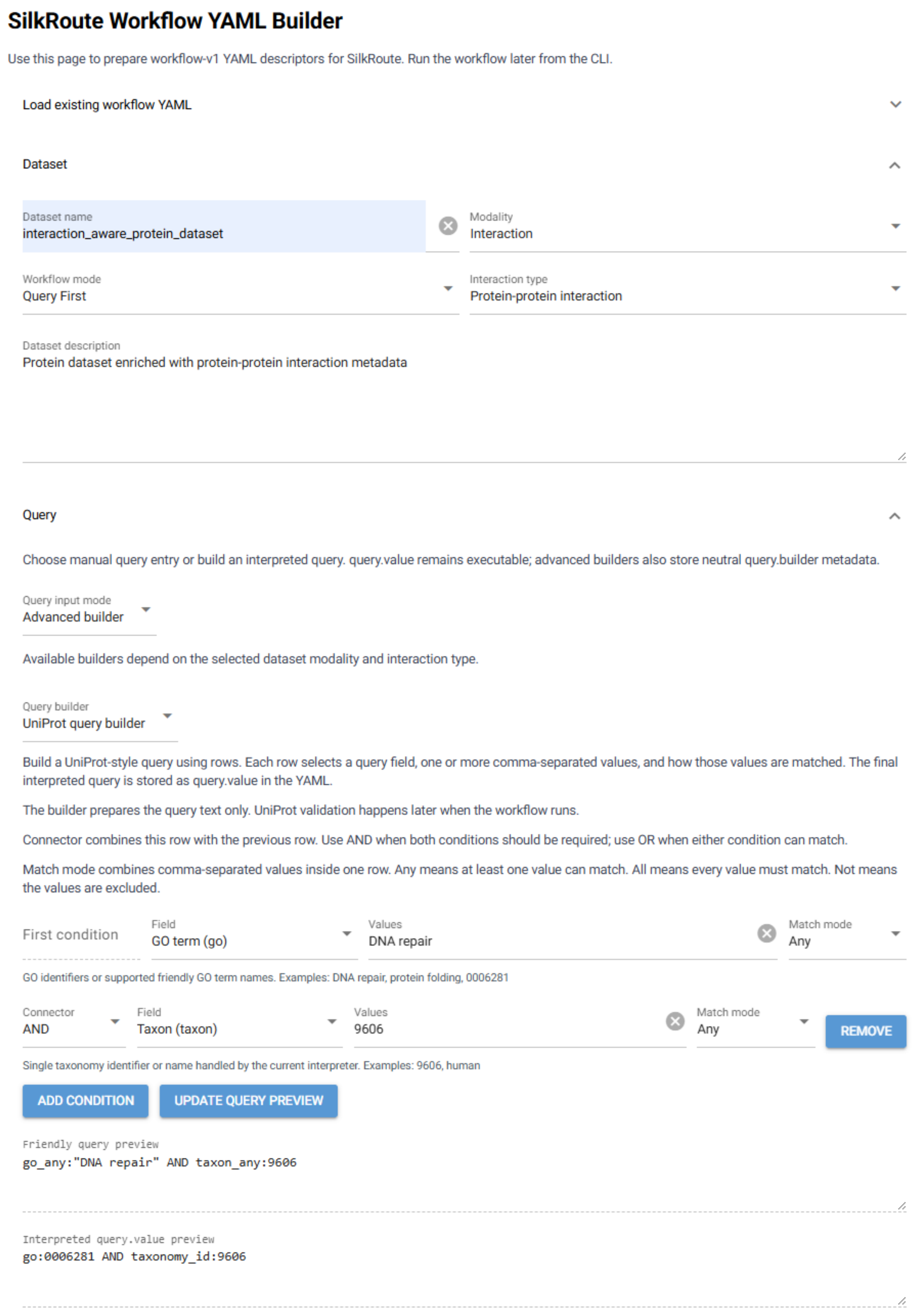

**Supplementary Figure S1**: Graphical interface for generating SilkRoute `workflow-v1` descriptors. The interface exposes the main descriptor sections, including dataset metadata, workflow mode, interaction type, and query configuration. The generated YAML file can be inspected, edited, versioned, and executed through the command-line workflow runner.

The descriptor defines the intended acquisition workflow. The following section describes how SilkRoute preserves output context, export structure, and execution metadata after the workflow is run.

### S4 Harmonization, Export, and Reproducibility Metadata

#### S4.1 Harmonization Descriptor Fields

SilkRoute exports primary and enrichment results as separate source-specific tables. Each interface retains the schema associated with its external resource rather than forcing heterogeneous outputs into a single universal representation. Consequently, the `harmonization` section should not be interpreted as a general cross-source schema-normalization stage.

The section contains optional fields that declare semantic column roles and provide limited export or reporting behavior. The `id_column` field specifies the name of a deterministic row-identifier column that is added to exported tabular copies when the requested column is not already present. Generated identifiers follow the output-specific row order and do not replace identifiers obtained from the original data source.

The `sequence_column` field identifies an existing column that can be inspected during reporting. When the named column is present in a top-level tabular output, SilkRoute counts its unique non-null values and records the result under `reporting.unique_sequences`. This field does not create, rename, normalize, or modify sequence values.

The `label_column` and `metadata_fields` entries are currently descriptive. They are validated and preserved in the workflow descriptor, `metadata.json`, and `run_summary.yml`, but they do not control column selection, renaming, merging, deduplication, or schema transformation. In particular, labels generated by `query_composition` workflows are written to the fixed `_label` column rather than to the column named by `label_column`.

A descriptor may record the intended semantic roles of its columns as follows:

```
harmonization:
id_column: record_id
label_column: activity_class
sequence_column: sequence
metadata_fields:
- accession
- protein_name
- organism_name
```

In this example, only `id_column` directly alters exported tabular copies, and `sequence_column` can affect generated reporting. The remaining entries document intended column roles without transforming the retrieved data. Omitting the complete `harmonization` section does not change query interpretation, source retrieval, enrichment, or the source-specific schemas returned by the workflow.

#### S4.2 Machine-readable Manifests and Reproducibility

SilkRoute records workflow configuration and execution results using both pre-execution descriptors and post-execution metadata files. This distinction is important because workflows that depend on external biological databases may produce different results over time as upstream resources are updated, corrected, expanded, or temporarily unavailable.

The YAML descriptor defines the intended workflow configuration before execution. It specifies the dataset modality, execution mode, query expression, requested fields, enrichment sources, runtime options, harmonization settings, and export behavior. This file is therefore the primary artifact for reviewing and re-running the intended acquisition procedure.

After execution, SilkRoute writes machine-readable files that describe what actually occurred during the run. The `metadata.json` file stores detailed execution metadata, including the original descriptor, normalized workflow values, source-specific parameters, execution status, output files, and available source-level metadata. The `run_summary.yml` file provides a more compact summary of the same run, including the dataset name, query, execution status, duration, output files, and record counts.

Table **S5** summarizes the role of these files.

**Supplementary Table S5**: Descriptor and metadata files produced or used by SilkRoute workflows.

| File | Role |
|---|---|
| `*.workflow-v1.yml` | User-provided descriptor that defines the intended workflow before execution. It should be treated as the authoritative workflow specification. |
| `metadata.json` | Detailed machine-readable execution manifest. It records the original descriptor, normalized workflow values, execution metadata, source-specific parameters, output files, and available enrichment metadata. |
| `run_summary.yml` | Compact execution summary intended for quick inspection. It reports execution status, runtime, output files, row counts, column counts, and selected workflow parameters. |
| `*.csv` | Tabular outputs generated by the workflow. These include the primary result table and any source-specific enrichment tables requested by the descriptor. |

This separation supports reproducibility and auditability at two levels. First, the YAML descriptor allows a workflow to be shared and re-executed. Second, the metadata and run summary files document the observed execution, including the exact normalized query parameters and the files generated during that run. This is especially useful when external services evolve over time or when a workflow is re-run in a different environment.

For example, a protein workflow may contain a user-facing query such as:

```
antimicrobial AND reviewed:true AND length:100-500
```

During execution, this query can be normalized into the syntax required by the target service. The descriptor preserves the original query written by the user, while the execution metadata records the normalized query submitted to the corresponding source interface. Together, these records make it possible to inspect both the intended query and the observed runtime behavior.

The metadata files should not be interpreted as replacing the exported tables. Instead, they provide context for interpreting them. The tables contain source-derived records, whereas `metadata.json` and `run_summary.yml` describe how those records were obtained, where they were written, and which workflow configuration produced them.

##### S4.3 Representative export schemas

SilkRoute workflows generate tabular data files together with machine-readable execution records. Exported dataset files contain the retrieved and parsed records, whereas execution metadata files describe the acquisition process, execution context, and generated outputs. The exact biological columns depend on the selected data source, endpoint, query, and enrichment settings; however, the export structure follows a common organization that separates dataset records from provenance and summary metadata.

**Supplementary Table S6**: Representative output files and schemas generated by SilkRoute workflows.

| **Output** | **Format** | **Purpose and representative content** |
|---|---|---|
| Dataset export | `csv`, `json`, `xml`, or `parquet` | Source-specific tabular output containing the retrieved or enriched records. When `harmonization.id_column` is configured and the requested column is absent, SilkRoute adds a deterministic row-identifier column to the exported copy. Other columns remain determined by the corresponding source interface, query fields, and output type. `metadata.json` |
| JSON | Workflow-level manifest describing the executed descriptor, selected data source, query snapshot, execution mode, enrichment settings, retry policy, source-specific methods or endpoints, fetched identifiers, failed identifiers, parsing summaries, execution times, and generated files. | |
| `run_summary.yml` | YAML | Compact execution summary intended for rapid inspection. It records the main workflow settings, output location, generated files, and high-level acquisition statistics when available. |
| Failed-record report | Text, CSV, JSON, or manifest field | Optional record of identifiers, queries, or records that could not be fetched, parsed, or exported. This output supports debugging and partial reruns. |
| Execution log | Console output or log file | Human-readable execution trace containing progress messages, warnings, source-specific errors, and debug information when enabled. |

The following case studies illustrate how these descriptor, export, and metadata mechanisms are used in representative SilkRoute workflows.

#### S5 Supplementary workflow examples and representative outputs

This section provides supplementary workflow configurations, execution examples, and representative outputs generated with SilkRoute. The first subsection corresponds to the antimicrobial protein case study described in the main text and is included to support reproducibility of the reported example. The remaining subsections provide additional workflow examples that illustrate other acquisition scenarios supported by the framework, including compound bioactivity retrieval from ChEMBL and interaction-aware protein-protein interaction dataset construction.

##### S5.1 Representative antimicrobial protein case study

This subsection provides the complete supplementary material associated with the representative antimicrobial protein dataset construction example described in the main text. It includes the YAML workflow descriptor, the command-line execution instruction, a programmatic Python example, and a summary of the generated output files. Together, these elements document the configuration, execution, and resulting files of the protein-centric `query_first` workflow used to retrieve reviewed antimicrobial protein records from UniProt and enrich them through AlphaFold, InterPro, PathwayCommons, and PDB cross-references.

##### S5.1.1 YAML workflow descriptor

The YAML dataset descriptor used for the antimicrobial protein case study is provided below. This configuration file defines the complete SilkRoute workflow used to retrieve reviewed antimicrobial protein records from UniProt, specify selected output fields, request cross-reference enrichment, and export the generated outputs together with execution metadata and summary files.

```yaml
schema_version: workflow-v1
dataset:
  name: antimicrobial_reviewed_proteins
  description: Reviewed UniProt protein records retrieved with an antimicrobial query.
  modality: protein
  mode: query_first
query:
  value: antimicrobial AND reviewed:true AND length:100-500
  include_isoform: false
  fields:
  - accession
  - protein_name
  - organism_name
  - sequence
  - organism_id
  crossref_fields:
  - alphafold
  - pdb
  - pathwaycommons_neighborhood
  - interpro
execution:
  enrich: true
  max_workers: 5
  total_retries: 3
  chembl_pages_to_fetch: -1
  debug: false
  download_alphafold_structures: false
  download_pdb_structures: false
harmonization:
  metadata_fields:
  - accession
  - protein_name
  - organism_name
  - sequence
export:
  output_dir: results/antimicrobial_reviewed_proteins
  format: csv
  graph_payload_storage: file
  graph_payload_compression: gzip
  include_metadata: true
  include_summary: true
  manifest_file: metadata.json
  summary_file: run_summary.yml
```

Listing 2: YAML dataset descriptor used for the antimicrobial protein case study.

##### S5.1.2 Command-line execution

The antimicrobial protein workflow was executed from the command line using the SilkRoute workflow runner. This route reproduces the complete descriptor-driven workflow, including descriptor validation, configured enrichment steps, output file generation, and creation of both `metadata.json` and `run_summary.yml`.

```
silkroute workflow run --config examples/workflows/antimicrobial_reviewed_proteins.workflow-v1.yml
```

Listing 3: execution command.

##### S5.1.3 Programmatic Python execution

SilkRoute can also be used programmatically through its public Python interface. The following example illustrates an in-memory protein query using the `Workflow` class.

```
from silkroute import Workflow

workflow = Workflow(
    query="antimicrobial AND reviewed:true AND length:100-500",
    modality="protein",
    mode="query_first",
    fields=[
        "accession",
        "protein_name",
        "organism_name",
        "sequence",
        "xref_alphafolddb",
        "xref_pdb",
        "xref_pathwaycommons",
        "xref_interpro",
        "organism_id",
    ],
    include_isoform=False,
)

result = workflow.run()
data = result.data
metadata = result.metadata
```

Listing 4: Programmatic protein query using the SilkRoute Python interface.

##### S5.1.4 Cross-reference enrichment

The primary UniProt table was subsequently used as input for the cross-reference enrichment layer. In this step, selected external endpoints were defined in the workflow descriptor and executed with `CrossRefEnricher`, allowing the UniProt-derived records to be complemented with additional annotations while preserving the traceability of each enrichment operation.

```
from silkroute.core.crossref_enricher import CrossRefEnricher, EndpointSpec

endpoint_specs = [
    EndpointSpec(database="alphafold", endpoint="prediction"),
    EndpointSpec(database="interpro", endpoint="entry"),
    EndpointSpec(
        database="pathwaycommons",
        endpoint="neighborhood",
        params={
            "datasource": ["reactome", "uniprot"],
            "pattern": ["interacts-with", "used-to-produce"],
            "subpw": True,
            "direction": "undirected",
            "limit": 1,
        },
    ),
    EndpointSpec(database="pdb", endpoint="entry"),
]

enricher = CrossRefEnricher(
    endpoint_specs=endpoint_specs,
    max_workers=5,
    total_retries=3,
)

enriched_data, enriched_metadata = enricher.enrich(
```

```
    data=uniprot_df,
    format="dataframe",
)

alphafold_df = enriched_data["alphafold_prediction"]
interpro_df = enriched_data["interpro_entry"]
pathwaycommons_df = enriched_data["pathwaycommons_neighborhood"]
pdb_df = enriched_data["pdb_entry"]
```

Listing 5: Programmatic cross-reference enrichment from a UniProt-derived table.

##### S5.1.5 Representative outputs

The antimicrobial protein workflow generated a set of tabular outputs and execution metadata files under the directory `results/antimicrobial_reviewed_proteins/`. The primary output corresponds to the UniProt-derived dataset, whereas the remaining files correspond to enrichment outputs generated from cross-references to AlphaFold, InterPro, PathwayCommons, and PDB. A summary of the representative output files is provided in Table **S7**.

**Supplementary Table S7**: Representative output files generated by the antimicrobial protein case study.

| Output file | Category | Dimensions | Description |
|---|---|---|---|
| `uniprot_results.csv` | Primary result | 2,444 × 24 | Reviewed antimicrobial protein records retrieved from UniProt. |
| `alphafold_prediction.csv` | Enrichment | 2,521 × 12 | AlphaFold metadata linked to retrieved UniProt entries. Local structure-file downloads were disabled. |
| `interpro_entry.csv` | Enrichment | 8,919 × 26 | InterPro entries, families, domains, signatures, and associated annotations when available. |
| `pathwaycommons_neighborhood.csv` | Enrichment | 123 × 11 | PathwayCommons neighborhood output represented as one row per source query, with external graph file references when available. |
| `pdb_entry.csv` | Enrichment | 9,139 × 12 | PDB entry metadata linked through UniProt cross-references. Local structure-file downloads were disabled. |
| `metadata.json` | Execution metadata | – | Detailed execution manifest containing the original descriptor, normalized workflow values, source-specific metadata, execution status, and output file information. |
| `run_summary.yml` | Execution summary | – | Compact summary of the workflow execution, including status, runtime, output files, and record counts. |

##### S5.1.6 PathwayCommons graph payload export

PathwayCommons neighborhood responses may contain large raw graph payloads. For this reason, the antimicrobial protein workflow used `graph_payload_storage: file` and `graph_payload_compression: gzip`. Under this configuration, the tabular output does not embed the raw `graph_json` payload directly in `pathwaycommons_neighborhood.csv`. Instead, it stores one row per source query with file references, compressed payload sizes, and SHA-256 checksums when non-empty graph payloads are available.

In this run, 123 non-empty PathwayCommons graph payloads were written as external compressed JSON files. No invalid or missing graph payload files were reported in the local analysis. A summary is provided in Table **S8**

**Supplementary Table S8**: PathwayCommons graph payload output for the antimicrobial protein case study.

| Item | Count |
|---|---|
| Rows in pathwaycommons_neighborhood.csv | 123 |
| External graph payload files | 123 |
| Rows without external graph files | 2,321 |
| Invalid or missing graph payload files | 0 |

The primary table, `uniprot_results.csv`, contains the UniProt records retrieved by the executable query. The enrichment tables preserve source-specific records from AlphaFold, InterPro, PathwayCommons, and PDB. These outputs are intentionally exported as separate files because each external source exposes different record structures and metadata fields.

##### S5.2 Additional workflow examples

The antimicrobial protein case study provides the most detailed example because it exercises primary retrieval, cross-reference enrichment, graph payload export, and execution metadata.

The following examples are included to illustrate additional SilkRoute workflow modes beyond the antimicrobial protein case study. These examples are not intended as independent biological analyses, but as representative execution scenarios showing how the framework handles compound activity retrieval, query composition, and interaction-aware protein dataset construction. Each example was executed using a YAML descriptor compliant with the `workflow-v1` schema and generated source-specific output tables together with execution metadata and a run summary.

###### S5.2.1 Compound bioactivity example: ChEMBL IC50 activity composition

The compound bioactivity example demonstrates the use of a query-composition strategy to retrieve ChEMBL activity records associated with the IC50 endpoint. The workflow defines multiple numeric IC50 intervals and assigns a descriptor-defined label to each interval, allowing the resulting records to be grouped according to the query component that retrieved them. Each IC50 macro is expanded into a structured ChEMBL activity query constrained by `standard_type = IC50`, `standard_units = nM`, and the corresponding `standard_value` comparisons.

###### S5.2.2 YAML dataset descriptors

In this workflow, the range expression IC50 is interpreted using strict lower and upper comparisons, corresponding to `standard_value > a` and `standard_value < b`. The three labeled intervals were therefore $0 < \text{IC50} < 10$ nM for `very_high_potency`, $10 < \text{IC50} < 100$ nM for `high_potency`, and $100 < \text{IC50} < 1000$ nM for `moderate_potency`. These intervals are mutually exclusive and exclude their endpoints. No records in the reported output had `standard_value` exactly equal to 0, 10, 100, or 1000 nM. The assigned label is propagated to the exported records through the `_label` column, preserving the link between each activity record and the query-composition component that produced it.

```yaml
schema_version: workflow-v1
dataset:
  name: chembl_ic50_activity_composition
  description: "Compound bioactivity dataset grouped by IC50 potency ranges using ChEMBL activity
      records.

   filtering_strategy: Query composition executes multiple labeled IC50 range queries. Each query
       is resolved as a ChEMBL activity search with standard_type equal to IC50 and standard_value
       constrained to the requested numeric range. The label assigned to each range is propagated
       to the exported records through the _label column."
  modality: compound
  mode: query_composition
query:
  value: ic50:0-10 AND standard_units:nM=very_high_potency,ic50:10-100 AND standard_units:nM=
      high_potency,ic50:100-1000 AND standard_units:nM=moderate_potency
  include_isoform: false
```

```yaml
  composition:
  - label: very_high_potency
    value: ic50:0-10 AND standard_units:nM
    builder:
      schema_version: query-builder-v1
      source: chembl
      builder_key: chembl_ic50_activity
      builder_type: ic50_activity
      rows:
      - comparison_mode: range
        lower_value: '0'
        upper_value: '10'
        value: ''
        standard_units: nM
  - label: high_potency
    value: ic50:10-100 AND standard_units:nM
    builder:
      schema_version: query-builder-v1
      source: chembl
      builder_key: chembl_ic50_activity
      builder_type: ic50_activity
      rows:
      - comparison_mode: range
        lower_value: '10'
        upper_value: '100'
        value: ''
        standard_units: nM
  - label: moderate_potency
    value: ic50:100-1000 AND standard_units:nM
    builder:
      schema_version: query-builder-v1
      source: chembl
      builder_key: chembl_ic50_activity
      builder_type: ic50_activity
      rows:
      - comparison_mode: range
        lower_value: '100'
        upper_value: '1000'
        value: ''
        standard_units: nM
execution:
  enrich: false
  max_workers: 5
  total_retries: 3
  chembl_pages_to_fetch: -1
  debug: false
export:
  output_dir: results/chembl_ic50_activity_composition
  format: csv
  include_metadata: true
  include_summary: true
  manifest_file: metadata.json
  summary_file: run_summary.yml
```

Listing 6: YAML dataset descriptor used for the ChEMBL IC50 activity composition case study.

##### S5.2.3 Representative outputs

The workflow generated a single ChEMBL output table containing 1,445,939 activity records and 47 columns. All retrieved records were associated with `standard_type = IC50` and `standard_units = nM`. The output contained 371,773 records labeled as `very_high_potency`, 494,205 records labeled as `high_potency`, and 579,961 records labeled as `moderate_potency`. The observed `standard_value` range was 0.0–999.82 nM, with a mean value of 176.79 nM.

**Supplementary Table S9**: Outputs generated by the ChEMBL IC50 activity composition workflow.

| Output file | Rows | Columns | Description |
|---|---|---|---|
| chembl_results.csv | 1,445,939 | 47 | Source-derived ChEMBL IC50 activity records with potency labels. |
| metadata.json | – | – | Workflow metadata, normalized descriptor, and ChEMBL retrieval metadata. |
| run_summary.yml | – | – | Compact execution summary and output file report. |

The purpose of this example is to show how SilkRoute can generate a labeled compound activity dataset from declarative range-based queries. The exported records remain source-derived activity records and are not further interpreted by SilkRoute. Downstream users can apply additional filtering, aggregation, compound standardization, duplicate handling, or target-specific selection according to the requirements of their analysis.

##### S5.2.4 Interaction-aware example: protein-protein interaction dataset

The interaction-aware example demonstrates the construction of a protein-protein interaction dataset starting from a biologically defined UniProt query. This workflow illustrates how SilkRoute can combine primary protein retrieval with interaction-oriented dataset generation, using a query-builder expression that captures both a functional criterion and a taxonomic constraint. In this case, the workflow targets human proteins associated with DNA repair and subsequently generates source-specific outputs suitable for downstream interaction analysis.

##### S5.2.5 YAML dataset descriptors

In this workflow, a structured query-builder expression was used to define an interaction-aware protein acquisition task. The query combined a functional condition, `DNA repair`, with the human taxonomy identifier `9606`. During workflow normalization, SilkRoute resolved the biological term `DNA repair` into the corresponding Gene Ontology identifier, generating the executable UniProt query `go:0006281 AND taxonomy_id:9606`. This preserves the link between the user-facing biological search intent and the normalized query used for reproducible dataset acquisition.

```yaml
schema_version: workflow-v1
dataset:
  name: interaction_aware_protein_dataset
  description: Protein dataset enriched with protein-protein interaction metadata
    from STRING and BioGRID.
  modality: interaction
  mode: query_first
  interaction_type: protein-protein
query:
  value: go:0006281 AND taxonomy_id:9606
  include_isoform: false
  builder:
    schema_version: query-builder-v1
    source: uniprot
    builder_key: uniprot
    builder_type: field_boolean
    rows:
    - connector: null
      field: go
      match_mode: any
      values:
      - DNA repair
    - connector: AND
      field: taxon
      match_mode: any
      values:
      - '9606'
```

```yaml
execution:
  enrich: false
  max_workers: 5
  total_retries: 3
  chembl_pages_to_fetch: -1
  debug: false
export:
  output_dir: results/interaction_aware_protein_dataset
  format: csv
  include_metadata: true
  include_summary: true
  manifest_file: metadata.json
  summary_file: run_summary.yml
```

Listing 7: YAML dataset descriptor used for the protein-protein interaction case study.

After the primary UniProt retrieval, the protein–protein interaction workflow queried the BioGRID `interactions` and STRING `interaction_partners` endpoints. BioGRID requests in the canonical execution were constructed from the `gene_primary` and `organism_id` values associated with each UniProt record, producing requests based on `geneList` and `taxId=9606`. Although the implementation can fall back to BioGRID identifiers when gene and organism information is unavailable, this fallback was not used by the interaction records in the canonical output.

The BioGRID interface used `start=0` and `max=10000`, with `interSpeciesExclude=false`, `selfInteractionsExclude=false`, `includeEvidence=false`, `searchBiogridIds=false`, and `searchIds=false`. The `evidenceList` parameter was omitted. Thus, the workflow did not apply an evidence-type filter and did not exclude self-interactions or interspecies interactions. The 10,000-record value represents a per-request maximum; no source query in the canonical execution reached that value, with the largest recorded response containing 6,108 interactions.

STRING requests used available `string_ids` when present. When a STRING identifier was unavailable, the query builder used `gene_primary` together with `species=9606`. The `interaction_partners` endpoint was queried with `network_type=functional`, while `required_score` and `limit` were omitted. Consequently, SilkRoute did not configure an explicit STRING confidence threshold or partner limit for this execution.

Both sources were parsed and exported as separate source-specific tables with provenance columns linking each interaction response to the corresponding primary UniProt query. SilkRoute performed no workflow-level interaction filtering, pair deduplication, or merge of BioGRID and STRING into a single interaction table.

###### S5.2.6 Representative outputs

The workflow generated a UniProt-derived protein table containing 2,253 records, together with source-specific interaction outputs from BioGRID and STRING. BioGRID produced 902,713 interaction records, while STRING produced 5,702 interaction partner records. These outputs demonstrate that SilkRoute can expand a biologically defined protein set into interaction-aware datasets while preserving separate files for each interaction source.

**Supplementary Table S10**: Outputs generated by the protein-protein interaction workflow.

| Output file | Rows | Columns | Description |
|---|---|---|---|
| `uniprot_results.csv` | 2,253 | 36 | UniProt proteins retrieved using the normalized DNA repair query. |
| `biogrid_interactions.csv` | 902,713 | 11 | BioGRID protein-protein interaction records linked to the retrieved proteins. |
| `string_interaction_partners.csv` | 5,702 | 18 | STRING functional interaction partner records. |
| `metadata.json` | – | – | Workflow metadata and interaction-source retrieval metadata. |
| `run_summary.yml` | – | – | Compact execution summary and output file report. |

The interaction records may include heterogeneous evidence types, repeated interaction pairs, source-specific identifiers, and different interaction confidence semantics. SilkRoute intentionally preserves these records as retrieved, allowing downstream users to decide how to filter, harmonize, deduplicate, or summarize the interaction evidence.

This example therefore illustrates two complementary aspects of the workflow system: users can define biologically meaningful query components through the query builder, while the executable query stored in the workflow remains explicit and reproducible. The resulting UniProt protein set was then expanded with protein-protein interaction records from BioGRID and STRING.

### S6 Supplementary Notes

#### S6.1 Current limitations

SilkRoute depends on external biological repositories, whose APIs, query syntax, pagination rules, authentication requirements, and response schemas may change over time. Although YAML descriptors, metadata files, and execution summaries preserve the executed configuration and provenance, they cannot guarantee that future repository queries will return identical records.

Another limitation is the heterogeneity and incompleteness of repository metadata. Different sources use distinct identifiers, evidence models, field names, and curation criteria. Therefore, harmonized outputs may still contain source-specific fields, missing values, or partially populated annotations. The absence of an annotation should not be interpreted as evidence that a biological property is absent.

Repository bias must also be considered. Public databases tend to overrepresent well-studied organisms, proteins, targets, diseases, and compounds, which may affect downstream analyses and machine learning models. Similarly, query-derived labels, such as activity ranges or disease-associated groups, should be interpreted as operational labels based on available annotations rather than definitive biological classes.

Finally, the construction of negative or unlabeled datasets remains challenging. Records not retrieved by a query should not automatically be treated as negatives, since they may correspond to untested or incompletely annotated cases. For this reason, supervised learning datasets generated from repository queries require additional validation and careful documentation of positive, negative, and unlabeled instances.

#### S6.2 Future developments

Future development will focus on extending SilkRoute as a more expressive and interoperable biomolecular dataset acquisition layer while preserving its current scope around reproducible retrieval, enrichment, provenance, and structured export. One direction is the incorporation of additional repositories and endpoint-specific modules, including resources that are currently only partially supported or not yet systematically integrated. Existing interfaces may also be expanded to cover additional retrieval and enrichment operations where these provide clear value for dataset-oriented acquisition workflows.

A second direction is the continued improvement of query construction and descriptor validation. Although the current implementation supports interpreted queries and predefined patterns for selected UniProt, ChEMBL, ChEBI, and PubChem searches, future versions could expose additional ontology-aware and preset-based query builders for organisms, reviewed status, sequence length, keywords, GO terms, EC numbers, temperature, pH, activity constraints, chemical names, and chemical structure patterns. Stricter validation of descriptor fields and source-specific query combinations could further detect unsupported or inconsistent configurations before execution. Natural language-assisted query construction may also be explored as a higher-level interface for translating user intent into explicit and validated repository-specific queries.

Future work will additionally address execution at larger scale, reporting, and interoperability. Planned directions include stronger support for containerized and distributed or cloud-native acquisition workflows, improved cache reuse, file checksums and release manifests, expanded validation of generated outputs, and more informative human-readable execution reports. FAIR-oriented metadata exports and integration with broader dataset-documentation standards and general workflow-execution systems may further improve the portability and archival value of SilkRoute acquisition packages. Additional representative acquisition scenarios, including peptide-, enzyme-, protein-engineering-, and compound-centered applications, will also be considered as the framework evolves.

##### S6.3 Code and Availability

SilkRoute is distributed as an open-source Python package under the MIT license. Source code, documentation, installation instructions, workflow descriptors, notebooks, and representative examples are provided through the project repository. The public package exposes a command-line entry point through `silkroute`, and the optional graphical interface can be installed through the GUI dependency group.

#### Supplementary References

[1] The UniProt Consortium. Uniprot: the universal protein knowledgebase in 2025. *Nucleic Acids Research*, 53(D1):D609–D617, 2025.

[2] Tamara Goldfarb, Vamsi K Kodali, Shashikant Pujar, Vyacheslav Brover, Barbara Robbertse, Catherine M Farrell, Dong-Ha Oh, Alexander Astashyn, Olga Ermolaeva, Diana Haddad, et al. Ncbi refseq: reference sequence standards through 25 years of curation and annotation. *Nucleic Acids Research*, 53(D1):D243–D257, 2025.

[3] Mihaly Varadi, Damian Bertoni, Paulyna Magana, Urmila Paramval, Ivanna Pidruchna, Malarvizhi Radhakrishnan, Maxim Tsenkov, Sreenath Nair, Milot Mirdita, Jingi Yeo, et al. Alphafold protein structure database in 2024: providing structure coverage for over 214 million protein sequences. *Nucleic Acids Research*, 52(D1):D368–D375, 2024.

[4] Helen M Berman and Stephen K Burley. Protein data bank (pdb): Fifty-three years young and having a transformative impact on science and society. *Quarterly Reviews of Biophysics*, 58:e9, 2025.

[5] Matthias Blum, Antonina Andreeva, Laise Cavalcanti Florentino, Sara Rocio Chuguransky, Tiago Grego, Emma Hobbs, Beatriz Lazaro Pinto, Ailsa Orr, Typhaine Paysan-Lafosse, Irina Ponamareva, et al. Interpro: the protein sequence classification resource in 2025. *Nucleic Acids Research*, 53(D1):D444–D456, 2025.

[6] Paul D Thomas, Dustin Ebert, Anushya Muruganujan, Tremayne Mushayahama, Laurent-Philippe Albou, and Huaiyu Mi. Panther: Making genome-scale phylogenetics accessible to all. *Protein Science*, 31(1):8–22, 2022.

[7] Gene Ontology Consortium. The gene ontology knowledgebase in 2026. *Nucleic Acids Research*, 54(D1):D1779–D1792, 2026.

[8] Igor Rodchenkov, Ozgun Babur, Augustin Luna, Bulent Arman Aksoy, Jeffrey V Wong, Dylan Fong, Max Franz, Metin Can Siper, Manfred Cheung, Michael Wrana, et al. Pathway commons 2019 update: integration, analysis and exploration of pathway data. *Nucleic Acids Research*, 48(D1):D489–D497, 2020.

[9] Marija Milacic, Deidre Beavers, Patrick Conley, Chuqiao Gong, Marc Gillespie, Johannes Griss, Robin Haw, Bijay Jassal, Lisa Matthews, Bruce May, et al. The reactome pathway knowledgebase 2024. *Nucleic Acids Research*, 52(D1):D672–D678, 2024.

[10] Minoru Kanehisa, Miho Furumichi, Yoko Sato, Yuriko Matsuura, and Mari Ishiguro-Watanabe. Kegg: biological systems database as a model of the real world. *Nucleic Acids Research*, 53(D1):D672–D677, 2025.

[11] Rose Oughtred, Jennifer Rust, Christie Chang, Bobby-Joe Breitkreutz, Chris Stark, Andrew Willems, Lorrie Boucher, Genie Leung, Nadine Kolas, Frederick Zhang, et al. The biogrid database: A comprehensive biomedical resource of curated protein, genetic, and chemical interactions. *Protein Science*, 30(1):187–200, 2021.

[12] Damian Szklarczyk, Katerina Nastou, Mikaela Koutrouli, Rebecca Kirsch, Farrokh Mehryary, Radja Hachilif, Dewei Hu, Matteo E Peluso, Qingyao Huang, Tao Fang, et al. The string database in 2025: protein networks with directionality of regulation. *Nucleic Acids Research*, 53(D1):D730–D737, 2025.

[13] Adnan Malik, Muhammad Arsalan, Carlos Moreno, Juan Mosquera, Eloy Félix, Tevfik Kizilören, Venkatesh Muthukrishnan, Barbara Zdrazil, Andrew R Leach, and Noel M O'Boyle. Chebi: re-engineered for a sustainable future. *Nucleic Acids Research*, 54(D1):D1768–D1778, 2026.

[14] Sunghwan Kim, Jie Chen, Tiejun Cheng, Asta Gindulyte, Jia He, Siqian He, Qingliang Li, Benjamin A Shoemaker, Paul A Thiessen, Bo Yu, et al. Pubchem 2025 update. *Nucleic Acids Research*, 53(D1):D1516–D1525, 2025.

[15] Barbara Zdrazil, Eloy Felix, Fiona Hunter, Emma J Manners, James Blackshaw, Sybilla Corbett, Marleen De Veij, Harris Ioannidis, David Mendez Lopez, Juan F Mosquera, et al. The chembl database in 2023: a drug discovery platform spanning multiple bioactivity data types and time periods. *Nucleic Acids Research*, 52(D1):D1180–D1192, 2024.

[16] Julia Hauenstein, Lisa Jeske, Antje Jäde, Mathias Krull, Katrin Dümmer, Julia Koblitz, Anja Tietz, Dieter Jahn, Lorenz Christian Reimer, and Boyke Bunk. Brenda in 2026: a global core biodata resource for functional enzyme and metabolic data within the dsmz digital diversity. *Nucleic Acids Research*, 54(D1):D527–D534, 2026.

[17] Ulrike Wittig, Maja Rey, Andreas Weidemann, Renate Kania, and Wolfgang Müller. Sabio-rk: an updated resource for manually curated biochemical reaction kinetics. *Nucleic Acids Research*, 46(D1):D656–D660, 2018.

[18] Parit Bansal, Anne Morgat, Kristian B Axelsen, Venkatesh Muthukrishnan, Elisabeth Coudert, Lucila Aimo, Nevila Hyka-Nouspikel, Elisabeth Gasteiger, Arnaud Kerhornou, Teresa Batista Neto, et al. Rhea, the reaction knowledgebase in 2022. *Nucleic Acids Research*, 50(D1):D693–D700, 2022.

[19] Uma Mudunuri, Anney Che, Ming Yi, and Robert M Stephens. biodbnet: the biological database network. *Bioinformatics*, 25(4):555–556, 2009.

[20] Yasset Perez-Riverol, Chakradhar Bandla, Deepti J Kundu, Selvakumar Kamatchinathan, Jingwen Bai, Suresh Hewapathirana, Nithu Sara John, Ananth Prakash, Mathias Walzer, Shengbo Wang, et al. The pride database at 20 years: 2025 update. *Nucleic Acids Research*, 53(D1):D543–D553, 2025.
